# Daedalus and Gasz recruit Armitage to mitochondria, bringing piRNA precursors to the biogenesis machinery

**DOI:** 10.1101/608018

**Authors:** Marzia Munafò, Vera Manelli, Federica A. Falconio, Ashley Sawle, Emma Kneuss, Evelyn L. Eastwood, Jun Wen Eugene Seah, Benjamin Czech, Gregory J. Hannon

**Affiliations:** Cancer Research UK Cambridge Institute, University of Cambridge, Li Ka Shing Centre, Cambridge CB2 0RE United Kingdom

## Abstract

The piRNA pathway is a small RNA-based immune system that silences mobile genetic elements in animal germlines. piRNA biogenesis requires a specialised machinery that converts long single-stranded precursors into small RNAs of ~25-nucleotides in length. This process involves factors that operate in two different subcellular compartments: the nuage/Yb-body and mitochondria. How these two sites communicate to achieve accurate substrate selection and efficient processing remains unclear. Here, we investigate a previously uncharacterized piRNA biogenesis factor, Daedalus (Daed), that is located on the outer mitochondrial membrane. Daed is essential for Zucchini-mediated piRNA production and for the correct localisation of the indispensable piRNA biogenesis factor, Armitage (Armi). We find that Gasz and Daed interact with each other and likely provide a mitochondrial “anchoring platform” to ensure that Armi is held in place, proximal to Zucchini, during piRNA processing. Our data suggest that Armi initially identifies piRNA precursors in nuage/Yb-bodies in a manner that depends upon Piwi and then moves to mitochondria to present precursors to the mitochondrial biogenesis machinery. These results represent a significant step in understanding a critical aspect of transposon silencing, namely how RNAs are chosen to instruct the piRNA machinery in the nature of its silencing targets.

## INTRODUCTION

The piRNA pathway acts in the germlines of animals as diverse as arthropods, amphibians, and mammals to control the expression of mobile genetic elements, protecting the genome from the potentially harmful consequences of uncontrolled transposon mobilisation (Czech et al. 2018). piRNAs function in complex with Argonaute proteins of the PIWI clade (in *Drosophila*: P-element induced wimpy testis (Piwi), Aubergine (Aub), and Argonaute-3 (Ago3)) guiding them to repress transposons at both transcriptional and post-transcriptional levels (Brennecke et al. 2007; Gunawardane et al. 2007; Sienski et al. 2012; Le Thomas et al. 2013; Rozhkov et al. 2013; Czech et al. 2018). This pathway has been extensively studied in *Drosophila*, where a number of genetic screens have uncovered many of its key components, some of which still await functional characterisation (Czech et al. 2013; Handler et al. 2013; Muerdter et al. 2013).

Animal germ cells harbour characteristic perinuclear structures that are required for the production of piRNAs. In *Drosophila* nurse cells these are called nuage and are the location wherein Aub/Ago3 ping-pong looping occurs (Brennecke et al. 2007; Gunawardane et al. 2007; Lim and Kai 2007; Malone et al. 2009). In follicle cells, piRNA precursors and biogenesis factors are concentrated in Yb-bodies, named after their main component, female sterile (1) Yb (Yb) (Szakmary et al. 2009; Olivieri et al. 2010; Saito et al. 2010; Qi et al. 2011; Murota et al. 2014). Germline piRNA biogenesis begins in nuage with the generation of 5’-monophosphorylated (5’-P) precursor RNAs via Aub/Ago3 slicing, a crucial event that specifies a cellular RNA as substrate for piRNA production (Han et al. 2015; Mohn et al. 2015; Senti et al. 2015; Wang et al. 2015; Gainetdinov et al. 2018). It is likely that a similar 5’-P precursor is generated without Aub nor Ago3 in Yb-bodies, but the underlying molecular mechanism for this process remains obscure. Following this initial precursor specification, the production of mature, Piwi-bound piRNAs occurs on the outer surface of mitochondria, where the conserved endonuclease Zucchini (Zuc) converts single-stranded 5’-P precursor RNAs into strings of consecutive piRNAs, each ~25-nucleotides (nt) in length (Ipsaro et al. 2012; Nishimasu et al. 2012; Han et al. 2015; Homolka et al. 2015; Mohn et al. 2015). During this process, binding of PIWI proteins to the 5’-P ends of the precursor RNAs is thought to help position Zuc, thus dictating the distinctive “phasing” of its cleavage (Gainetdinov et al. 2018). In essence, the PIWI footprint on the nascent piRNA precursor determines the 5’ end of the next piRNA in this processive cycle (Brennecke et al. 2007). Interestingly, the mitochondrial localisation of the piRNA biogenesis machinery is generally conserved across species, strongly implying a functional role for mitochondria in piRNA biology and transposon defence.

Several other piRNA biogenesis factors are also localized to mitochondria, including the Tudor domain-containing, <u>Pa</u>rtner of <u>PI</u>WIs (Papi), the glycerol-3-phosphate acyltransferase, Minotaur (Mino), and the <u>G</u>erm cell protein with <u>A</u>nkyrin repeats, <u>S</u>terile alpha motif, and leucine <u>Z</u>ipper (Gasz) (Liu et al. 2011; Czech et al. 2013; Handler et al. 2013; Vagin et al. 2013; Hayashi et al. 2016). With the exception of Papi, which is largely dispensable in flies but involved in piRNA 3’ formation in other species (Honda et al. 2013; Hayashi et al. 2016; Nishida et al. 2018), loss of any of these factors severely impairs Zuc-mediated piRNA generation. Compromised mitochondrial piRNA biogenesis results in Piwi proteins lacking bound piRNAs, which are consequently destabilised and degraded, ultimately leading to the transcriptional de-repression of transposons (Wang and Elgin 2011; Sienski et al. 2012; Le Thomas et al. 2013; Rozhkov et al. 2013). In contrast, Aub/Ago3-mediated slicing of precursors and the ping-pong cycle are unaffected by loss of these factors.

Besides mitochondrially localized proteins, a number of cytosolic factors contribute to the process of piRNA biogenesis. Among these is Armitage (Armi), an RNA helicase of the Upf1 family, which localises to nuage and mitochondria in germ cells and predominantly to Yb-bodies in follicle cells (Malone et al. 2009; Olivieri et al. 2010; Saito et al. 2010). Armi shows ATP-dependent 5’-3’ helicase activity (Pandey et al. 2017) and Zuc-mediated piRNA biogenesis, but not the ping-pong cycle, collapses upon its loss. Tethering of Armi to a reporter transcript results in conversion of the RNA into ~25-nt piRNAs (Pandey et al. 2017; Rogers et al. 2017). The mouse homolog of Armi, MOV10L1, also binds to piRNA precursors and initiates the production of piRNAs (Vourekas et al. 2015). As a whole, these data place Armi at a critical juncture in piRNA biogenesis, where its binding to precursor transcripts is both necessary and sufficient to specify downstream piRNA production by Zuc and its mitochondrial cofactors.

Our current model of piRNA biogenesis identifies two subcellular compartments as being critical for piRNA production, nuage/Yb-bodies, where precursor transcripts are recognised and processed into 5’-P intermediates, and mitochondria, where such intermediates are processively cleaved into mature piRNAs. While these two structures are often in physical proximity, it is unclear how they specifically interact to promote piRNA biogenesis. Here we identify a novel piRNA biogenesis factor, CG10880/Daedalus (Daed), which is anchored on the mitochondrial outer membrane. We show that Daed, together with Gasz, provides a mitochondrial binding platform for Armi, which is, in turn, essential for Zuc-mediated production of piRNAs. Our data suggests that Armi moves from nuage/Yb-bodies, where it associates with piRNA precursors and Piwi, to mitochondria, where it remains in close association with Zuc during the processive cycle of piRNA production. We propose that loss of Gasz or Daed leads to impaired production of piRNAs due to the inability of the Armi-Piwi complex to be stably recruited to the mitochondrial surface where it delivers precursor RNAs to Zuc.

## RESULTS

### CG10880/Daedalus is a mitochondrially localized protein required for piRNA biogenesis

Comprehensive genetic screens in *Drosophila* have provided a molecular parts list for the piRNA pathway (Czech et al. 2013; Handler et al. 2013; Muerdter et al. 2013), yet how a number of these factors act to promote piRNA production or transposon silencing remains to be understood. Among such factors was CG10880, an uncharacterized *Drosophila* protein required for transposon silencing in the germline compartment of the ovary (Czech et al. 2013).

The gene encoding CG10880 is located on the left arm of chromosome 2 and shows its highest expression in ovarian tissues (Fig.S1A). CG10880 contains a Sterile Alpha Motif (SAM, often involved in protein-protein or protein-RNA interaction), a coiled-coil domain (CC, typically involved in protein oligomerisation and linked to diverse cellular functions), and a predicted transmembrane domain (TMM) at its carboxyl-terminus (Fig.1A). Depletion of *CG10880* from the fly germline resulted in transposon de-repression at levels comparable to those observed for knockdowns of *zuc* and *gasz* (Fig.S1B) and in a strong delocalisation of Piwi from nuclei, a hallmark of impaired piRNA biogenesis (Fig.S1C). Interestingly, the CG10880 domain structure resembles that of Gasz (Fig.S1D), a mitochondrial protein involved in piRNA biogenesis that also carries a SAM domain and a TMM at its carboxyl-terminus.

**Figure 1.**
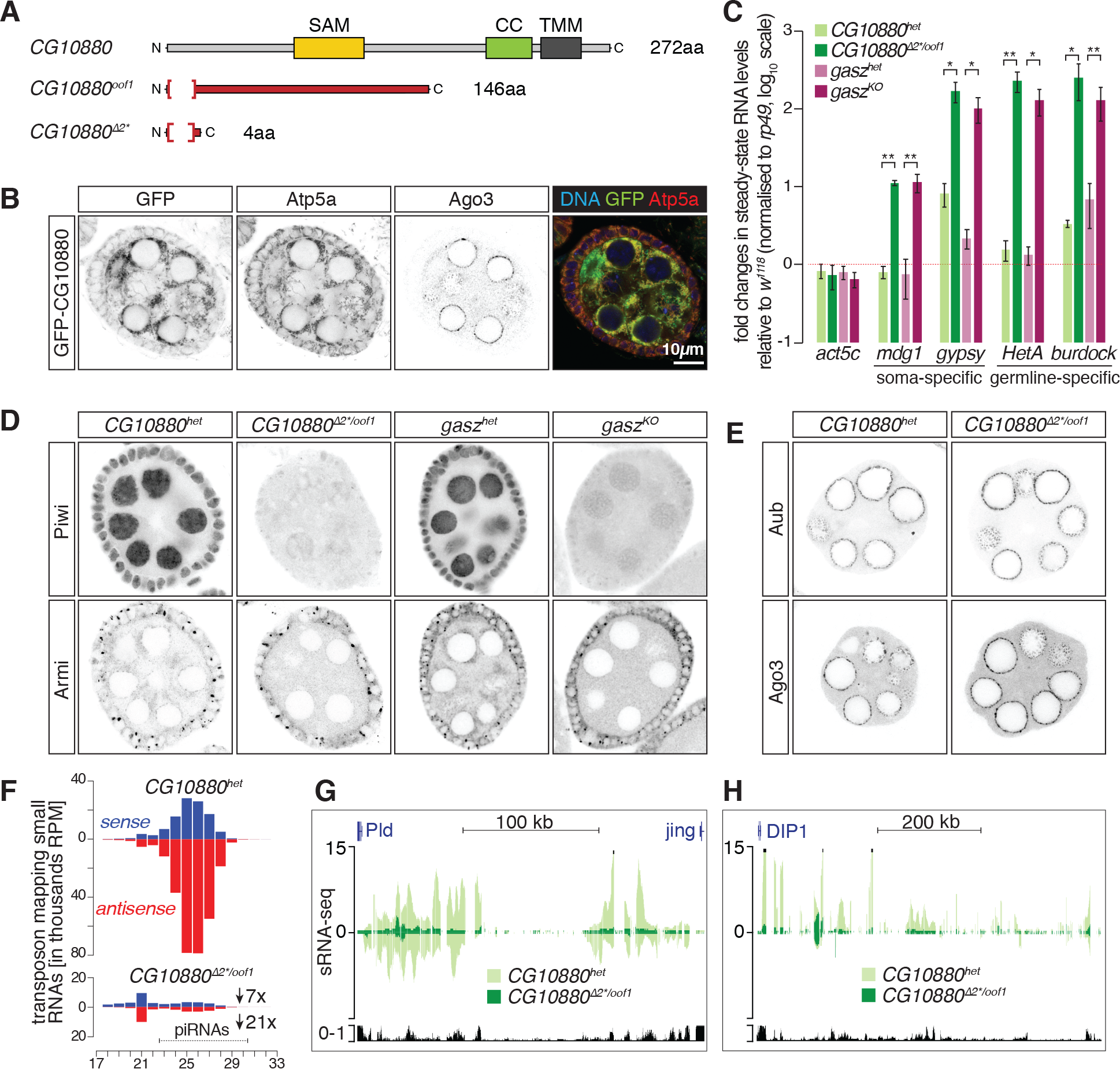
Daedalus is a mitochondrially localized protein involved in piRNA biogenesis. **(A)** Schematic representation of the CG10880/Daedalus domain structure and of the two null alleles. SAM = Sterile Alpha Motif, CC = Coiled Coil, TMM = transmembrane domain. Highlighted in red is the portion of coding sequence that is out of frame in the mutants. **(B)** Confocal images of GFP-CG10880 in ovaries (see also Supplementary Fig.S1E). Scale bar, 10µm. **(C)** Fold changes in steady-state RNA levels of the indicated soma- and germline-specific transposons from ovaries. Values are relative to *w^1118^* flies and normalised to *rp49*. * = P value < 0.05; ** = P < 0.001 (unpaired t-test). Error bars indicate standard deviation (n=4). **(D-E)** Confocal images of Piwi, Armi, Aub and Ago3 in ovaries (see also Supplementary Fig.S2C-E). **(F)** Size distribution of transposon-mapping small RNAs from ovaries. Sense reads are shown in blue, antisense in red. **(G-H)** Coverage plots of small RNA reads uniquely mapped to the dual-strand cluster *42AB* (**G**) and the uni-strand cluster *flamenco* (*flam*) (**H**). Shown are normalised reads per million (RPM). The mappability for an average 25 bp read length is shown at the bottom.

To determine the localisation of CG10880 in the fly ovary, we generated a line ubiquitously expressing an amino-terminally GFP-tagged fusion protein, GFP-CG10880, thus preserving its putative transmembrane domain. GFP-CG10880 localisation overlapped with the mitochondrial marker Atp5a and was adjacent to but separate from nuage, as marked by Ago3 (Fig.1B). Zuc-GFP (ubiquitously expressed) and GFP-Gasz (expressed from its endogenous promoter; described in (Handler et al. 2013)) showed an indistinguishable localisation pattern (Fig.S1E). These results suggested that CG10880 could function as a mitochondrial piRNA biogenesis factor and prompted us to generate CG10880 null mutants. We derived two alleles, one harbouring a deletion causing a frameshift (*CG10880^oof1^*) and a second deletion that results in a premature stop codon (*CG10880^∆2*^*) (Fig.1A). Homozygous mutant females laid fewer eggs, which showed abnormal morphology without dorsal appendages and did not hatch, similarly to *gasz* homozygous mutants (Fig.S2A), generated via RFP knock-in in the gasz genomic locus (*gasz^KO^*, Fig.S2B). *CG10880* trans-heterozygous and *gasz* homozygous mutants displayed impaired repression of both somatic and germline transposons (Fig.1C). Piwi nuclear localisation was severely compromised in somatic and germline cells of mutant flies (Fig.1D and S2C), whereas the nuage localisation of Aub, Ago3 and Vasa was unperturbed (Fig.1E and S2D). This implied that CG10880 is likely involved in the generation of Piwi-loaded piRNAs but not the ping-pong cycle. Interestingly, *CG10880* mutants also displayed an altered distribution of the RNA helicase Armi, which is normally localised to nuage and mitochondria (Fig.1D and S2C). Notably, the same phenotype is observed in *gasz* mutant (Fig.1D and S2C) and knockdown flies (Handler et al. 2013). Finally, *CG10880* null mutants had highly altered mitochondrial morphology (Fig.S2E), again closely resembling what is observed upon *gasz* loss (Fig.S2E and (Handler et al. 2013)). These results suggest that CG10880 is a bona-fide piRNA biogenesis factor involved in Zuc-mediated processing of phased piRNAs on mitochondria and, since it is essential for the correct assembly of the mitochondrial “labyrinth” in germ cells, we name it Daedalus (Daed).

In germ cells, piRNA biogenesis is initiated in nuage where the ping-pong cycle, driven by Aub and Ago3, generates long 5’-P piRNA precursors. These are further processed by Zuc on the outer mitochondrial surface resulting in the sequential generation of phased piRNAs. With the goal of understanding which biogenesis step is affected by the loss of Daed, we sequenced small RNAs from mutant ovaries. Repeat-derived small RNAs were dramatically reduced in *daed* trans-heterozygous mutants, as compared to their heterozygous siblings (6.9- and 20.8-fold for sense and antisense, respectively; Fig.1F), whereas repeat-derived siRNAs remained unchanged (21-nt peak in Fig.1F). piRNAs originating from germline dual-strand clusters, somatic uni-strand clusters, and protein-coding genes were all strongly reduced (Fig.1G,H and S3A), indicating an essential role of Daed in Zuc-dependent processing of piRNA precursors in both major ovarian cell types. Consistent with this hypothesis, the ping-pong signature of repeat-derived piRNAs was unaffected (Fig.S3B). As expected, small RNAs in *gasz^KO^* recapitulated the phenotype of *daed* mutants (Fig.S3B-C-D).

### Daedalus is essential for recruitment of Armi to mitochondria

Ovarian somatic cells (OSCs), cultured *in vitro*, express a functional Piwi-piRNA pathway (without the ping-pong cycle) and therefore provide a convenient context in which to investigate piRNA biogenesis (Niki et al. 2006; Saito et al. 2009). Immunostaining of OSCs transfected with 3×FLAG-tagged Daed showed localization to mitochondria, but removal of the putative TMM domain caused its redistribution throughout the cell (Daed^∆TMM^) (Fig.2A). Additionally, Daed co-localises with both Zuc and Gasz (Fig.2A).

**Figure 2.**
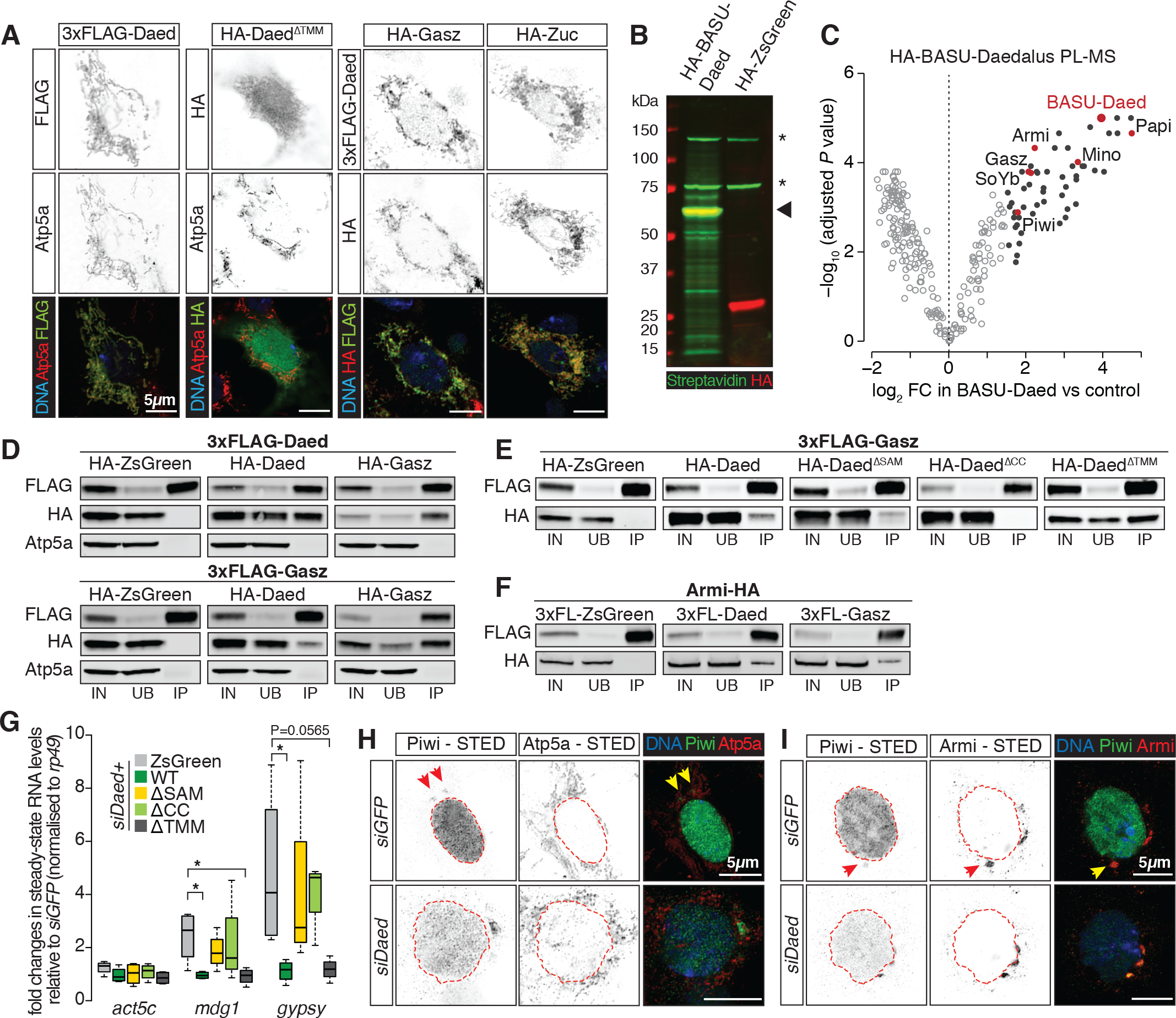
Daedalus interacts with Gasz and these together promote Armi localization on mitochondria. **(A)** Confocal images of fusion constructs and the mitochondrial marker Atp5a in OSCs. Scale bar, 5µm. **(B)** Western blot showing biotinylated proteins (in green, detected with Streptavidin) upon expression of HA-BASU-Daed compared to the HA-ZsGreen control. Asterisks indicate endogenously biotinylated proteins, the arrowhead indicates the size of HA-BASU-Daed fusion. Note that HA-BASU-Daed fusion biotinylates itself (green and red signal overlap). **(C)** Volcano plot showing enrichment and corresponding significance of biotinylated proteins identified via quantitative Mass Spectrometry from OSCs expressing BASU-Daed versus control (n=2). Black dots indicate proteins showing a log_2_FC > 1.5 and adjusted P value < 0.05 in BASU-Daed. Highlighted in red are piRNA pathway factors. Full list of enriched proteins is provided in **Table S1**. **(D-F)** Western blots of FLAG-tag co-immunoprecipitation from lysates of S2 cells transfected with the indicated constructs (IN, input; UB, unbound; IP, immunoprecipitate). **(G)** Fold changes in the steady-state RNA levels of somatic transposons in OSCs nucleofected with siRNAs and various rescue constructs. Values are relative to *GFP* control knockdown and normalised to *rp49*. * = P value < 0.05; ** = P < 0.001 (unpaired t-test) (n=4). **(H-I)** Stimulated Emission Depletion (STED) microscopy of Piwi and Atp5a **(H)** or Piwi and Armi **(I)** in OSCs from indicated knockdowns. Scale bar, 5µm.

Aiming to understand the role of Daed in piRNA biogenesis, we set out to identify its interacting partners via *in vivo* proximity labelling with biotin (Roux et al. 2012; Kim et al. 2016). The strength and stability of the biotin-Streptavidin interaction allows very stringent pulldown conditions that successfully isolate membrane proteins and, as a further advantage, proximity labelling allows the capture of even weak or transient interactions that would escape detection with standard IP-MS techniques. We found that the <u>*Ba*</u>*cillus* <u>*su*</u>*btilis* biotin ligase (BASU; (Ramanathan et al. 2018)) showed robust activity in fly cells at 26°C, and therefore expressed an HA-BASU-Daed fusion in OSCs. Western blot on lysates of cells expressing the fusion showed the appearance of biotinylated proteins in addition to those endogenously present (marked by asterisks in Fig.2B), when compared to cells expressing ZsGreen without biotin ligase. Biotinylated proteins were efficiently recovered using Streptavidin beads (Fig.S4A) and subjected to quantitative Mass Spectrometry (from now on referred to as Proximity Labelling-Mass Spectrometry, PL-MS). BASU-Daed itself was highly enriched in the pulldown (consistent with the self-biotinylation of any BASU fusion protein, indicated by arrowhead in Fig.2B), along with other known piRNA pathway factors (Fig.2C and **Table S1**). Strikingly, Daed PL-MS enriched for mitochondrial (Papi, Mino, and Gasz) as well as cytosolic (SoYb, Piwi and Armi) piRNA pathway proteins. Similarly, BASU-Gasz PL-MS also enriched for Armi and Piwi (Fig.S4B and **Table S2**). Together with the observation that Armi is mislocalised in *gasz* and *daed* mutants, these data raise the possibility that a Gasz/Daed transmembrane complex might anchor Armi onto mitochondria to achieve efficient piRNA production.

To test this hypothesis, we co-expressed Daed, Gasz and ZsGreen in S2 cells and probed their interaction by anti-FLAG co-immunoprecipitation. Both 3×FLAG-Daed and 3×FLAG-Gasz co-immunoprecipitated with HA-Daed and HA-Gasz but not with HA-ZsGreen, suggesting both a homo- and hetero-typic interactions on the mitochondrial surface (Fig.2D), though we cannot rule out the possibility that other proteins act as bridges in a larger complex. The mitochondrial marker Atp5a showed no enrichment in the immunoprecipitation, implying that the association of Gasz and Daed is not an artefact of intact mitochondria being isolated (Fig.2D). To further investigate Daed and Gasz homo- or heterotypic interaction, we expressed 3×FLAG-tagged Daed, Gasz, or Zuc in S2 cells. The latter served as a positive control, since Zuc is known to exist in a dimeric conformation (Ipsaro et al. 2012; Nishimasu et al. 2012). We chemically crosslinked the cells to stabilise any putative complexes and performed anti-FLAG pulldowns. Upon crosslinking, Zuc immunoprecipitation showed a second band at double the size of the fusion protein itself, thus likely corresponding to the dimer (light green arrowhead in Fig.S4C). Strikingly, a similar pattern was observed for Gasz and Daed (purple and dark green arrowhead in Fig.S4C), but not for ZsGreen.

To identify the regions that mediate these interactions, we performed 3×FLAG-Gasz co-immunoprecipitation with Daed deletion constructs lacking individual domains (Fig.S4D). These experiments revealed that the Daed-Gasz interaction depends on the CC domain of Daed (Fig.2E). Considered together, these data indicate that Gasz and Daed interact on the mitochondrial surface as direct binding partners. HA-tagged Armi also co-immunoprecipitated with both Gasz and Daed, thus indicating that its enrichment in Daed and Gasz PL-MS reflects interactions within the same complex, rather than just physical proximity (Fig.2F).

Knockdown of *daed* in OSCs leads to somatic transposon de-repression (Fig.S4E), which could be rescued by re-expression of siRNA-resistant Daed^WT^ and Daed^∆TMM^, but not those lacking the SAM or CC domains (Fig.2G). Daed^∆TMM^ could still interact with mitochondrial Gasz and function normally, in contrast with Daed^∆SAM^ which, albeit still associating with Gasz, appeared to be unable to exert its role. This potentially implicates the SAM domain in the interaction with either Armi or RNA. Piwi nuclear localisation was markedly reduced in OSCs depleted of Daed (Fig.S4F), and Piwi appeared to be retained with Armi in Yb-bodies (arrow in Fig.S4F), a phenotype that was also observed in follicle cells of *daed* and *gasz* mutant flies (arrows in Fig.S4G). These results suggest that, in the absence of Daed and Gasz, Piwi and Armi fail to leave Yb-bodies to translocate to mitochondria. We therefore exploited high resolution imaging using <u>St</u>imulated <u>e</u>mission <u>d</u>epletion (STED) microscopy to better understand the consequences of Daed depletion in OSCs. In wild-type cells, Piwi was detected in close association with mitochondria (arrows in Fig.2H, Fig.S4H), whereas upon *daed* knockdown the majority of the remaining Piwi became confined to discrete Yb-bodies, surrounded by morphologically altered mitochondria (Fig.2H). Co-staining of Piwi and Armi showed that, when outside of the nucleus, Piwi was generally observed in proximity with Armi (Fig.2I, S4I). Taken together, these data suggest a model in which Piwi moves onto the mitochondrial surface, where Armi is positioned in a manner dependent on Daed and Gasz, but is unable to reach these processing sites upon depletion of *daed* or *gasz*.

### Armi shuttles from Yb-bodies to mitochondria where it associates with dimeric Zuc

To gain a better understanding of protein-protein interactions occurring on the mitochondrial surface during piRNA biogenesis, we also carried out PL-MS for Armi and Zuc (Fig.3A,B and S5A,B). Armi-BASU PL-MS enriched for Yb and Piwi (Fig.3A), both previously reported as Armi interactors (Olivieri et al. 2010; Saito et al. 2010), thus validating the sensitivity of our method. In addition, we noted enrichment of other cytosolic (Shu, SoYb, Spn-E) and mitochondrial (Papi, Gasz, Daed, Mino) piRNA biogenesis factors (Fig.3A and **Table S3**). Interestingly, PL-MS for Zuc-BASU identified several mitochondrial components of the piRNA biogenesis machinery (Papi, Mino, Daed, Gasz), but also demonstrated strong enrichment of Armi and, to a lesser extent, SoYb (Fig.3B and **Table S4**), implying tight association of these factors during piRNA production. Structural studies strongly indicate that Zuc cleaves RNA as a dimer (Ipsaro et al. 2012; Nishimasu et al. 2012), thus we decided to exploit this feature to refine further our proteomics analysis and attempt to pinpoint which piRNA pathway factor is more closely associated with the “cleavage-competent” Zuc dimer. We applied the Split-BioID method in which the *E.coli* biotin ligase BirA* is divided into two fragments (N- and C-terminus BirA*), each inactive on its own (Schopp et al. 2017) (Fig.S5C, lanes 2-3). N- and C-BirA* reconstitute the active enzyme only if fused to two proteins that interact *in vivo*, as in the case of the Zuc dimer, and upon biotin supplementation (Fig.S5C, last lane, and S5D). The reconstituted BirA* exhibited much lower activity than BASU, thus generating a smaller number of biotinylated proteins (Fig.S5D). Strikingly, PL-MS using Split-BioID revealed enrichment of a limited set of proteins, yet still readily identified Armi (Fig.3C and **Table S4**). This could imply a closer association of Armi with the Zuc dimer than, for instance, SoYb, which was only identified by Zuc-BASU PL-MS. Knockdown of *zuc* and *armi* in OSCs caused more similar changes in the levels of genome-mapped small RNAs (r^2^=0.782, Fig.S5E), than *zuc* versus *yb* depletion (r^2^=0.406, Fig.S5F), further supporting a role for Armi as a proximate Zuc co-factor.

**Figure 3.**
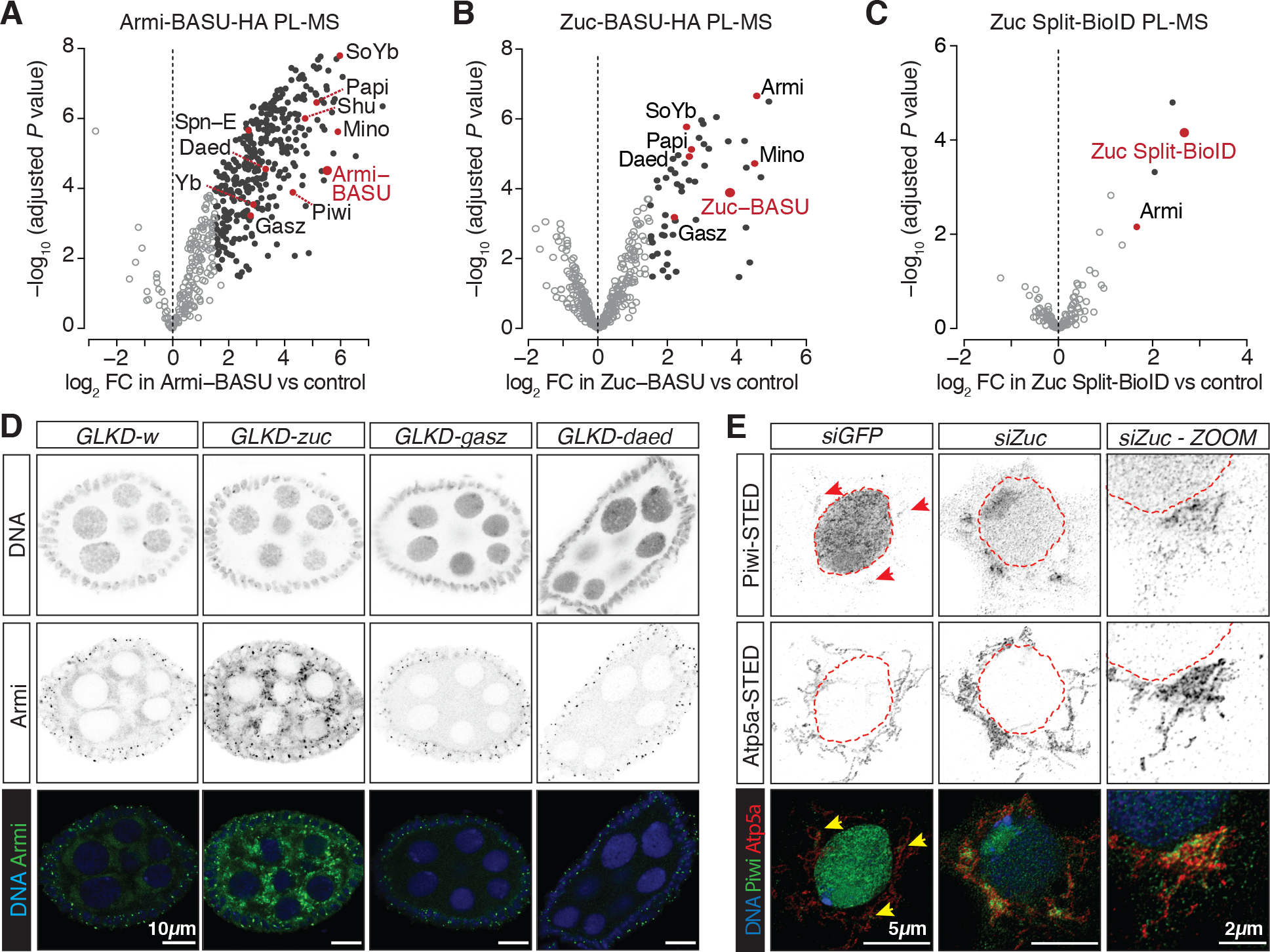
Armi localises to mitochondria in proximity to the piRNA processing machinery. **(A-C)** Volcano plots showing enrichment and corresponding significance of biotinylated proteins identified via PL-MS from OSCs expressing the indicated constructs against control (n=3). Black dots indicate proteins showing a log_2_FC > 1.5 and adjusted P value < 0.05. Highlighted in red are piRNA pathway factors. Full list of enriched proteins is provided in **Tables S3-5**. **(D)** Confocal images of Armi in ovaries from indicated GLKD. Scale bar, 10µm. **(E)** STED microscopy of Piwi and Atp5a in OSCs. Scale bar, 5µm.

Based upon this, we envisioned a model in which a Piwi-Armi complex is licensed at the sites of pre-piRNA specification (nuage in germ cells and Yb-body in follicle cells/OSCs) and then translocates to mitochondria. There, Armi is held in place by Gasz/Daed and engages in piRNA production in close association with Zuc. Consistent with this model, in both fly germline and OSCs, loss of *zuc* causes a dramatic accumulation of Piwi and Armi on mitochondria, whereas loss of *gasz* and *daed* lead to their dispersal in the cytosol or concentration in Yb-bodies (Fig.3D-E and Fig.S5G). We therefore suggest that Armi shuttles from nuage/Yb-bodies to mitochondria and is involved in the presentation of piRNA precursors to Zuc enabling their downstream processing into phased piRNAs.

### Armi depends upon Piwi for binding to piRNA precursors

Armi belongs to the family of Upf1-like RNA helicases and its mouse homolog MOV10L1 has been shown to bind to pre-piRNAs (Vourekas et al. 2015). Therefore, we sought to determine whether *Drosophila* Armi associates with the piRNA precursors that will be presented to Zuc for phased cleavage. CLIP-seq for an Armi-HALO fusion expressed in OSCs (Fig.S6A) showed substantial enrichment of somatic piRNA source transcripts: the uni-strand piRNA clusters *flamenco* (*flam*) and *20A* (red in Fig.4A) and a number of protein-coding genes known to give rise to genic piRNAs (shown in blue in Fig.4A). We found Armi distributed along the entire length of piRNA precursor transcripts, even when they span several hundred kilobases, such as in the case of *flam* (Fig.4B). On genic transcripts, Armi crosslinked preferentially to their 3’UTRs, as exemplified by *tj* (Fig.4C). In all cases, sequences enriched in Armi CLIP-seq corresponded with those appearing as piRNAs that were lost upon *armi* knockdown (Fig.4B-C, bottom panels). When analysing the presence of transposon content in Armi CLIP-seq, we found an enrichment for antisense sequences, especially those that are present in *flam* (red in Fig.4D, with dot size proportional to their abundance within *flam*). We did not detect substantial enrichment of transposon sense sequences (Fig.4E). No 1U bias was detected in Armi CLIP-seq, but this is likely a result of our library preparation procedure and is also consistent with what is reported for mouse MOV10L1 (Vourekas et al. 2015). Thus, our data supports a model in which Armi specifically binds to a subset of cellular transcripts and assists their processing into piRNAs. However, a key issue remains as to how such precursors are selectively discriminated by Armi from other cellular RNAs.

**Figure 4.**
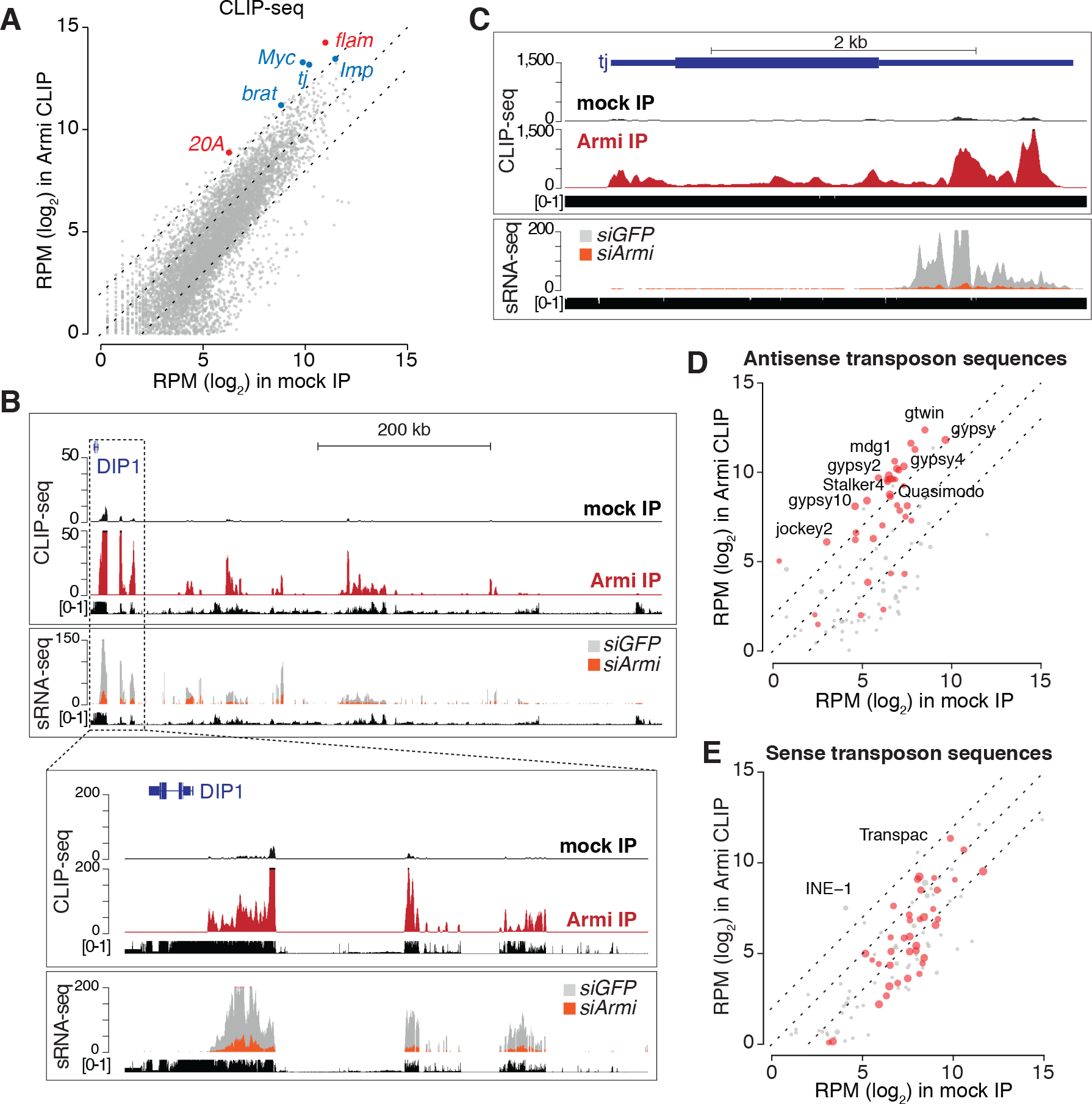
Armi binds to piRNA precursors. **(A)** Scatter plot showing expression levels (RPM) of genes in Armi CLIP-seq (n=4) against a mock IP (n=3). piRNA clusters expressed in OSCs are highlighted in red, selected protein coding genes producing piRNAs in blue. **(B)** UCSC genome browser shot displaying Armi CLIP-seq and small RNA-seq reads uniquely mapping to the piRNA cluster *flam* (upper panel) and a zoomed-in view of the first ~50 kb (bottom panel). Shown are normalised RPM. The mappability tracks for 50 bp and 25 bp read length, respectively, are shown below. **(C)** Same as in (B) but showing the protein-coding gene *tj*. **(D-E)** Scatter plots showing expression levels (RPM) of antisense and sense transposon sequences in Armi CLIP-seq against a mock IP. Transposon sequences present in *flam* are highlighted in red with dot size proportional to their abundance within *flam* according to dm6 Repeat Masker annotations.

The prevailing model suggests that Zuc simultaneously forms piRNA 3’ and 5’ ends by cleaving downstream of Piwi, while Piwi is positioned on the 5’ end of a longer piRNA precursor (Gainetdinov et al. 2018). Precisely how Armi fits into this process remains unclear, yet it does definitively also engage piRNA precursors. We therefore examined the interactions between Armi and piRNA precursors in the context of either *piwi*, *zuc* or *gasz* knockdown (Fig.5A-C, Fig.S6C-E). Upon depletion of Piwi, we detected a substantial decrease, but not complete loss, of the binding of Armi to *tj*, *flam*, and *20A*, the main sources of piRNAs in OSCs (Fig.5A). We suggest that the binding is not entirely lost due to the persistence of some Piwi protein in knockdown samples (Fig.S6B). In contrast, upon knockdown of *zuc*, Armi CLIP-seq indicated an increase in precursor transcript binding (Fig.5B). Interestingly, the Armi footprint on the *tj* mRNA was not evenly affected by *zuc* depletion, but importantly, increases were restricted to the 3’UTR, which is precisely the part of the *tj* mRNA that is converted into piRNAs. Finally, *gasz* knockdown did not globally affect Armi CLIP-seq signal (Fig.5C), which is in accordance with our model postulating that Armi binds to precursor RNAs before translocating on mitochondria. Quantification of CLIP-seq signals for selected regions of *flam* and *20A* that show good mappability (orange boxes in Fig.5A-C) and for *tj* CDS, 5’UTR, and 3’ UTR confirmed increased association of Armi with precursors in *zuc* knockdowns, while *piwi* depletion resulted in reduced Armi binding (Fig.5D).

To investigate whether the dependency of Armi precursor binding upon Piwi might stem from a physical association between these proteins, we immunoprecipitated Armi-3×FLAG and probed for the presence of endogenous Piwi. Immunoprecipitation of Armi-3×FLAG from wild type cells resulted in the recovery of only a small amount of Piwi (Fig.5E, quantification in Fig.S6F). However, upon *zuc* knockdown, despite an overall reduction in Piwi levels, the amount of Piwi complexed with Armi rose, and this association was insensitive to RNase (Fig.5E, S6F). Interestingly, *gasz* and *daed* knockdown did not generally impact the association of Armi and Piwi but instead made that interaction sensitive to RNase treatment (Fig.5E, S6F).

**Figure 5.**
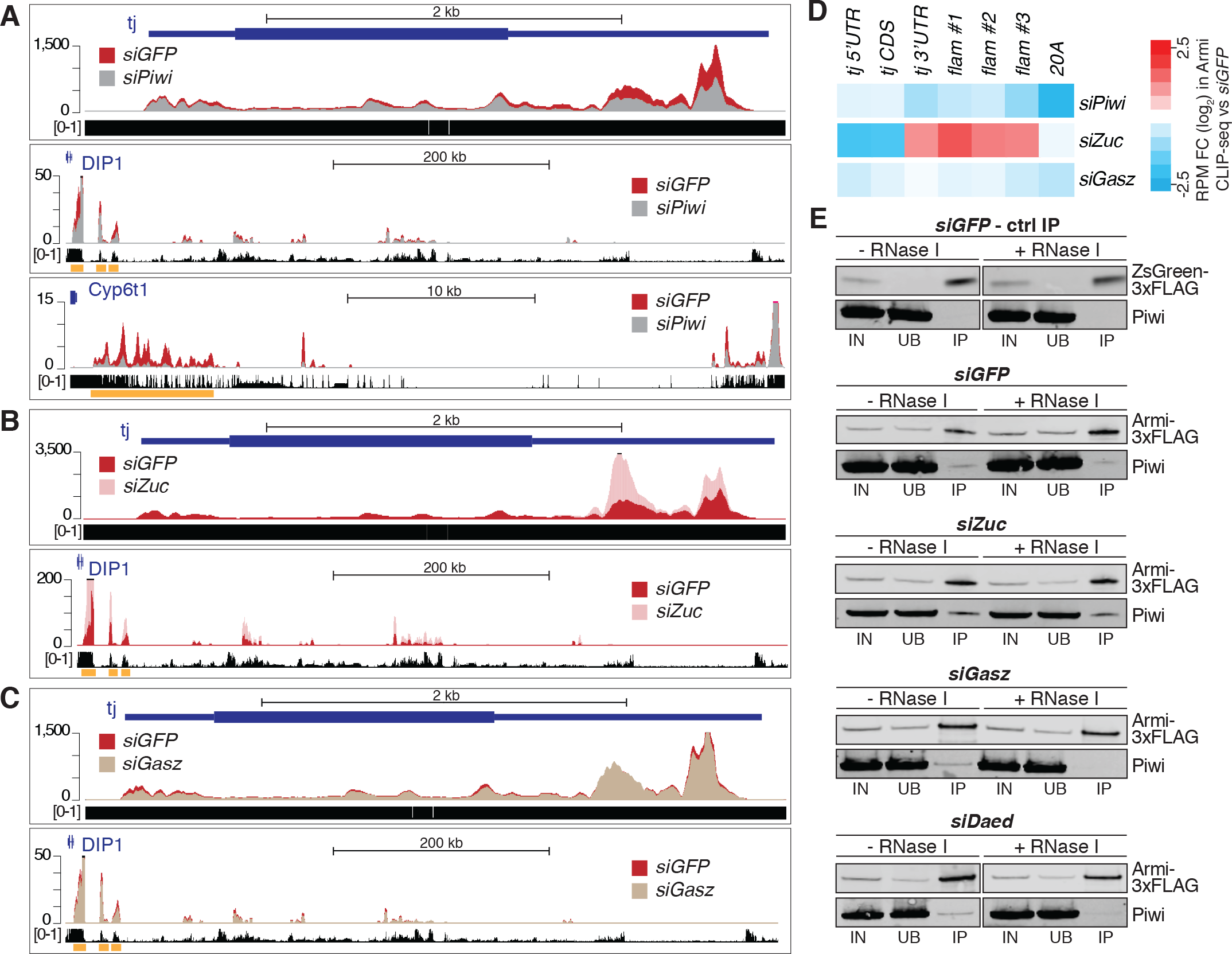
Armi binding to piRNA precursors is impaired upon Piwi knockdown. **(A-C)** Genome browser shot displaying Armi CLIP-seq profiles from OSCs upon indicated knockdown. Shown are normalised RPM. (n=3 for *siZuc* and relative *siGFP* control; n=4 for *siPiwi* and relative *siGFP* control; n=3 for *siGasz* and relative *siGFP* control). Below each profile are the mappability tracks for 50 bp read length. **(D)** RPM Log_2_ Fold-Change in Armi CLIPseq upon indicated knockdown against their relative *siGFP* for selected regions of *tj, flam* and *20A* (orange boxes in panels **A-C). (E)** Western blots of ZsGreen- or Armi-3×FLAG co-immunoprecipitation from lysates of OSCs upon indicated knockdown (IN, input; UB, unbound; IP, immunoprecipitate).

## DISCUSSION

The biogenesis of piRNAs requires a highly specialised machinery that must recognise the correct precursors in Yb-bodies or nuage, transport them to the surface of mitochondria and parse them into trails of ~25-nt piRNAs. How each step is achieved and how information flows between the discrete subcellular compartments in which piRNA biogenesis is initiated and completed is yet to be fully understood.

Here, we expand the repertoire of mitochondrial piRNA biogenesis factors by identifying and characterising CG10880/Daedalus (Daed). Daed is predominantly expressed in the female germline and appears to be unique to *Drosophilids* (Fig.S7). Its domain structure is similar to that of Gasz, a previously described mitochondrial piRNA biogenesis factor that is conserved across species. Although it is not clear why *Drosophilids* possess two proteins with related structure and function, our data indicate that Daed and Gasz assemble as homo- and hetero-polymeric complexes (Fig.6, middle) and act together to promote localisation of Armi on mitochondria in a non-redundant manner. It is likely that recruitment of Armi to the mitochondrial surface is key for delivery of piRNA precursor transcripts to the nuclease, Zuc, and the importance of Daed/Gasz in this process is confirmed not only by our own data but also by the conservation of a role for Gasz in transposon control across animals (Zhang et al. 2016). Intriguingly, Gasz and Daed loss similarly perturbs mitochondrial morphology, and Zuc depletion has been shown to result in mitochondrial clustering (Olivieri et al. 2012). It is presently unclear how changes in piRNA biogenesis produce such dramatic morphological impacts, but this observation further underscores the intimate relationship between mitochondria and the transposon control machinery in the ovary.

**Figure 6.**
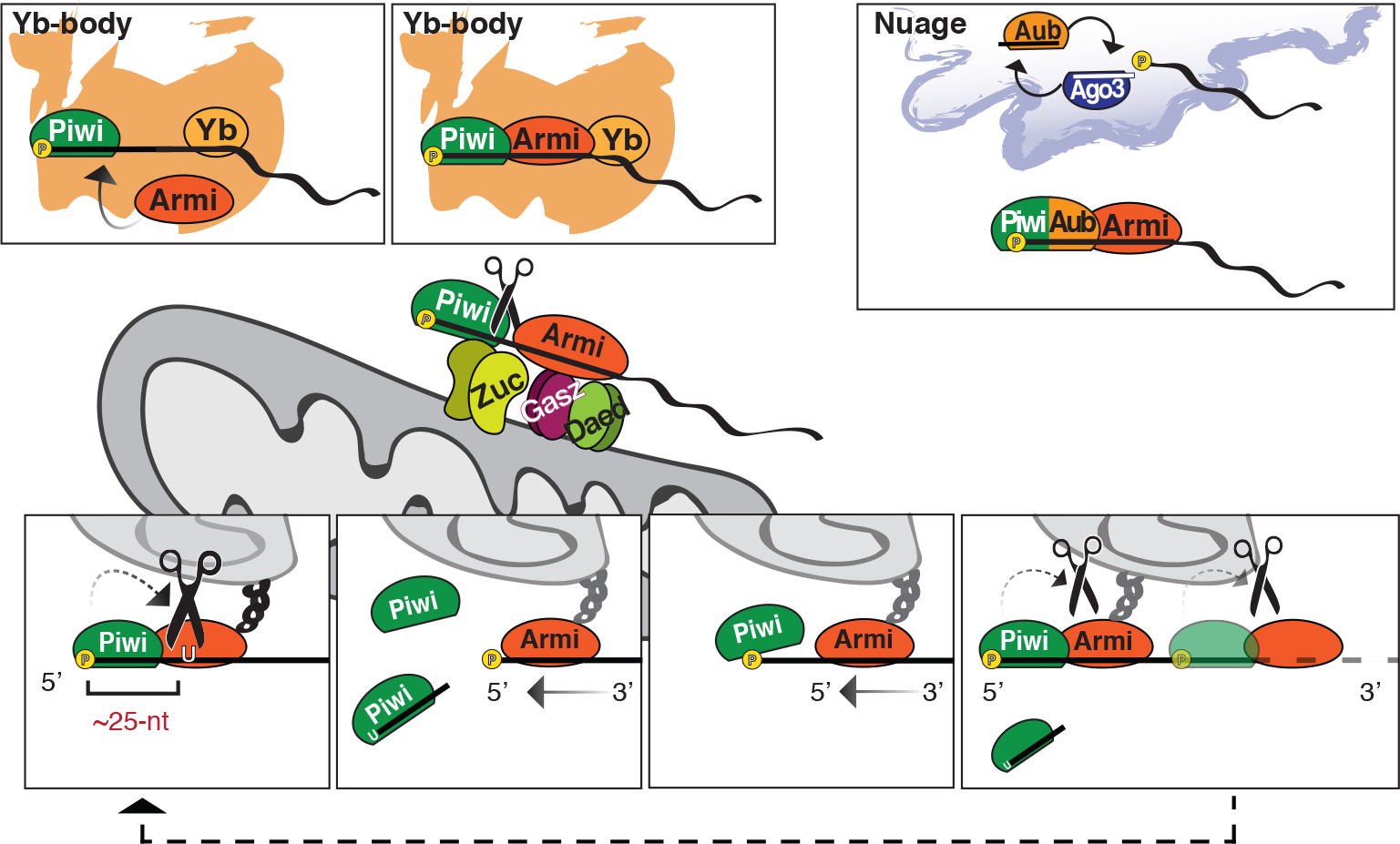
A model for piRNA biogenesis. In Yb-bodies, Piwi binds to the 5’ phosphate of a piRNA precursor, and this in turn recruits Armi [top]. Piwi, Armi, and the substrate RNA translocate to mitochondria where a Daed/Gasz complex stabilizes Armi while Zuc dimers cleave the transcript [middle]. Phased piRNA production then requires cycles of Piwi binding positioning Zuc to cleave at the first available uridine. While held in place by Daed and Gasz, Armi translocates along the transcript to provide the next segment of the RNA for piRNA biogenesis, and again Piwi binding to the newly-generated 5’-P restarts the process [bottom].

Our data indicate that the sites of piRNA precursor specification and processing communicate via translocation of precursors marked by binding of Piwi and Armi from Yb-bodies/nuage to dimeric Zuc on the mitochondrial surface. Though our proteomics experiments identified several cytosolic piRNA biogenesis factors as being adjacent to mitochondria, among those Armi appeared to be the one in closest proximity to Zuc. This could imply that Armi’s role is to ensure the processivity of Zuc cleavage, by presentation of RNA substrates via its ATP-dependent RNA helicase activity. Indeed, upon *zuc* knockdown, Armi and Piwi are trapped on mitochondria in an RNA-bound state, presumably because the subsequent step in piRNA generation is blocked. It is interesting to note that Armi binding to RNA depended on Piwi (Fig.5A). This provides a potential link between the recognition of a piRNA precursor in nuage and Yb-bodies, via Piwi/Aub binding to its 5’-P end, and the subsequent association with Armi and flow into downstream processing. Since all factors involved are expressed in both compartments of the fly ovary, this model might equally apply to nurse and follicle cells.

Considered as a whole, our data support a model (Fig.6) whereby 5’-P piRNA precursors in nuage or Yb-bodies are first bound by Piwi or Aub. Substrates defined in this way can then recruit Armi. This complex must then translocate to mitochondria where Gasz and Daed anchor Armi adjacent to dimeric and active Zuc, potentially via its associated precursor RNA. Once held in place on the mitochondrial surface, the Armi-Piwi interaction is stabilised independently of RNA, and the cycle of piRNA production can initiate. At this stage, mitochondrially anchored Armi likely unwinds or translocates along the precursor RNAs to allow Piwi to sequentially bind to each free 5’-P end, generated after each Zuc cleavage event. The Piwi footprint in turn determines the next Zuc cleavage site upstream of the first accessible uridine. Yet, what mechanism dictates this particular Zuc cleavage preference remains an outstanding question.

---

**Figure S1.**
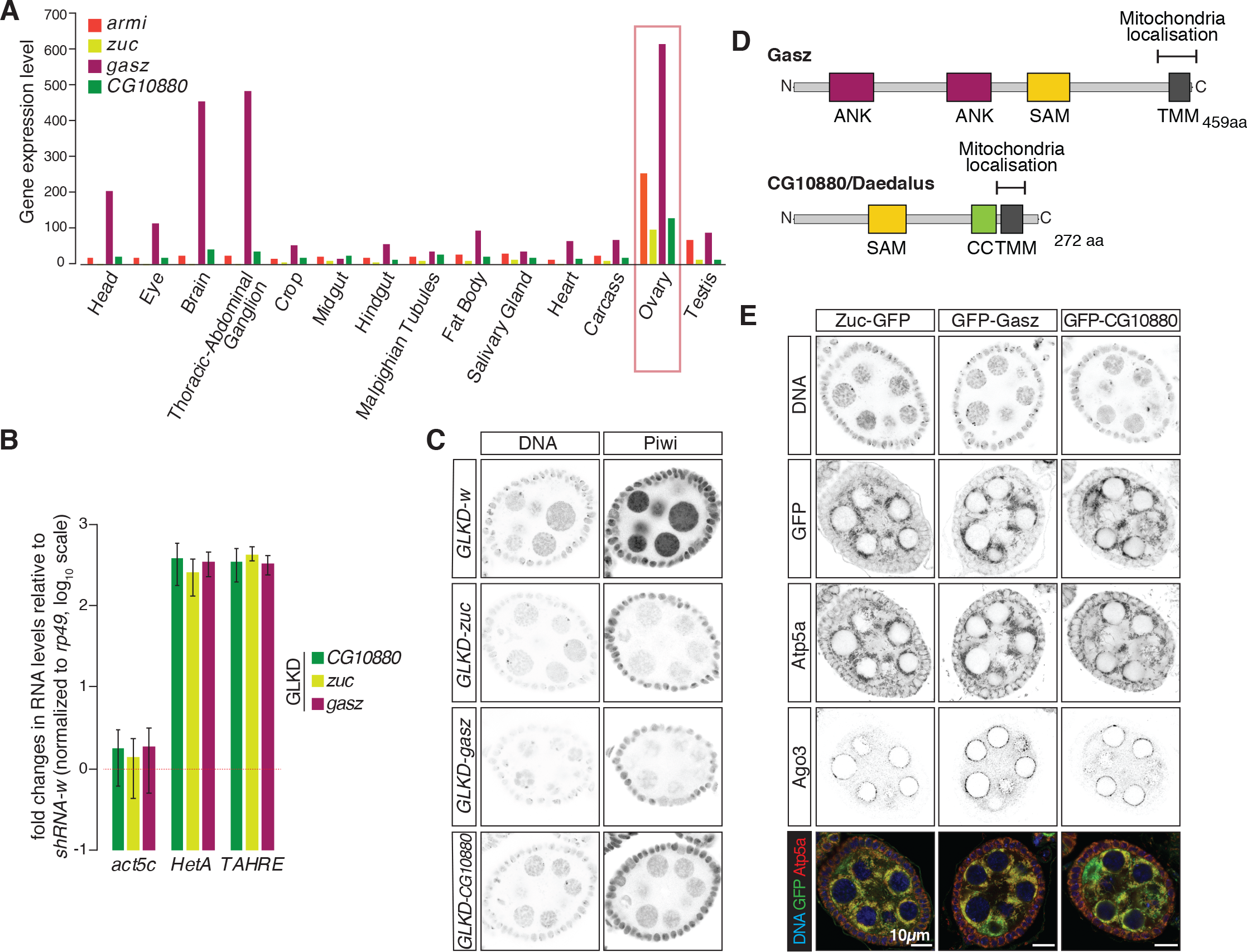
**(A)** Expression levels of *armi*, *zuc*, *gasz* and *CG10880* in various tissues of the adult fly. **(B)** Fold changes in the steady-state RNA levels of transposons from GLKD ovaries. Values are relative to *w* knockdown and normalised to *rp49*. Error bars indicate standard deviation (n=2). **(C)** Confocal images of Piwi in ovaries upon indicated GLKD. Scale bar, 10µm. **(D)** Schematic representation of the Gasz and CG10880/Daedalus domain structure. ANK = Ankyrin repeats, SAM = Sterile Alpha Motif, CC = Coiled Coil, TMM = transmembrane domain **(E)** Confocal images of Zuc-GFP, GFP-Gasz and GFP-CG10880 in ovaries. Scale bar, 10µm.

**Figure S2.**
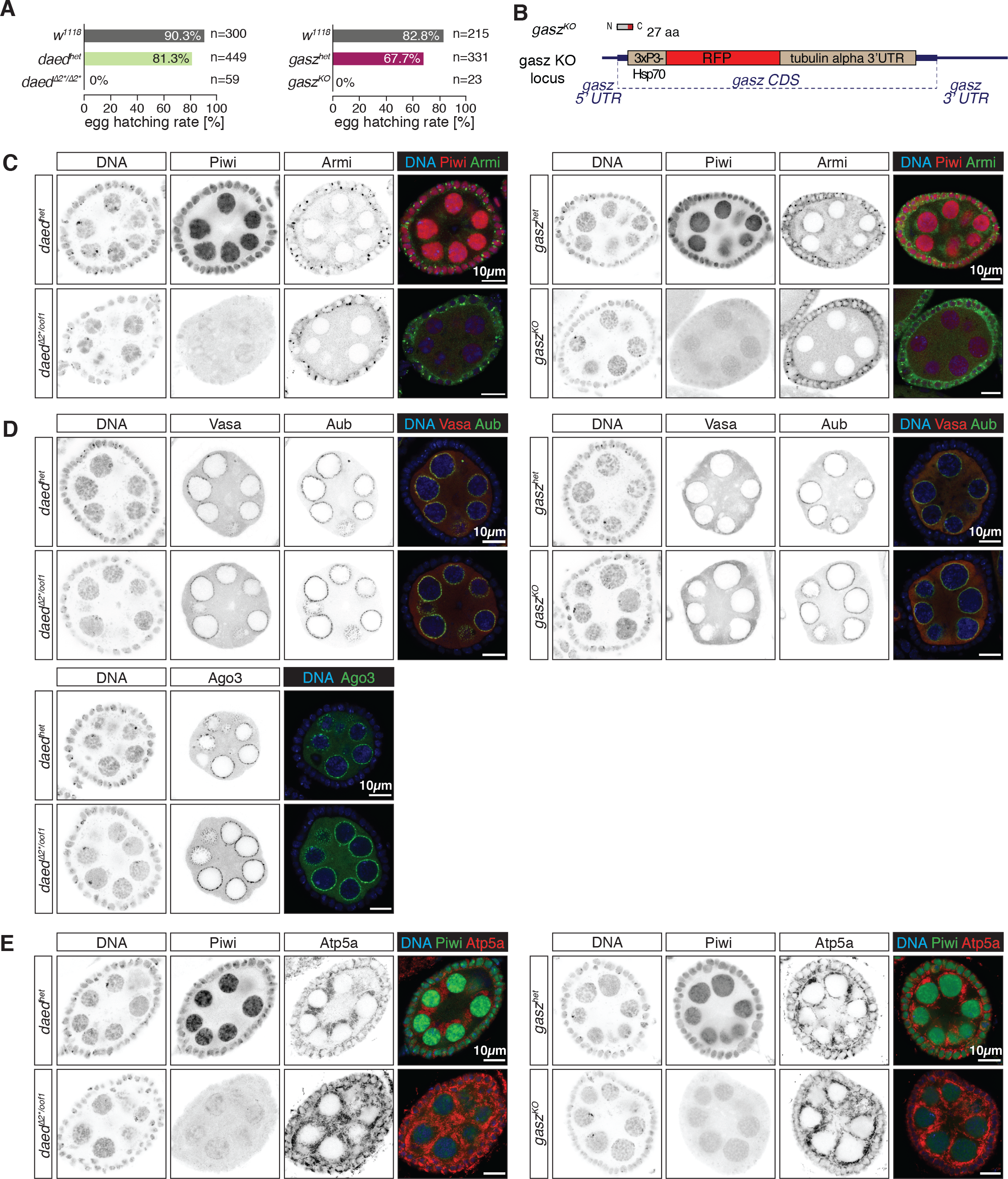
**(A)** Bar graphs showing the egg hatching rates of female flies of the indicated genotypes. **(B)** Schematic representation of the gasz^KO^ allele. **(C-E)** Confocal images of Piwi, Armi, Vasa, Aub, Ago3 and Atp5a. Scale bars, 10µm.

**Figure S3.**
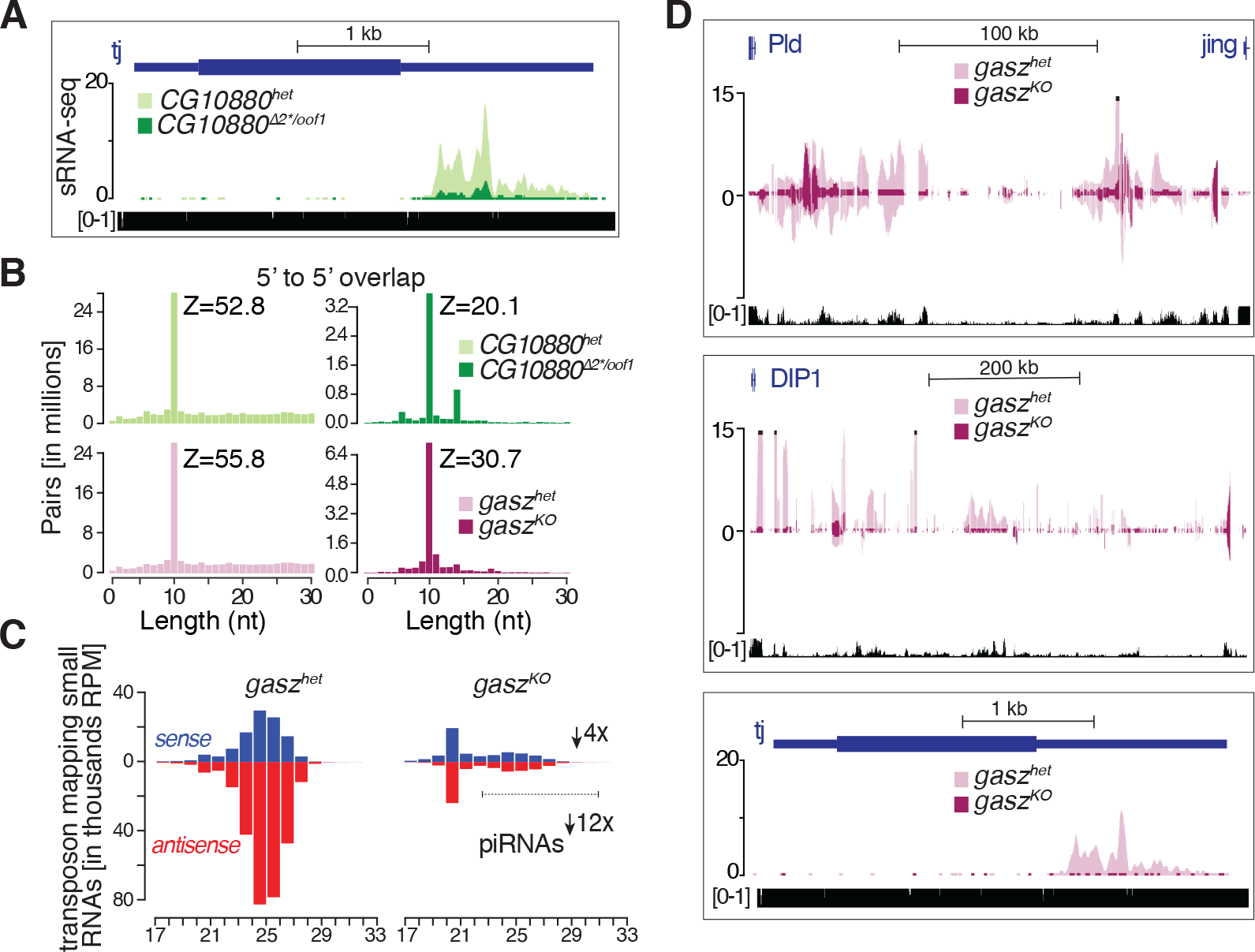
**(A,D)** Coverage plots of uniquely mapped small RNA reads from ovaries. Shown are normalised RPM. The mappability, calculated for an average 25 bp read length, is shown at the bottom. **(B)** Ping-pong analysis of transposon-mapping small RNAs from ovaries. **(C)** Size distribution of transposon-mapping small RNAs from ovaries.

**Figure S4.**
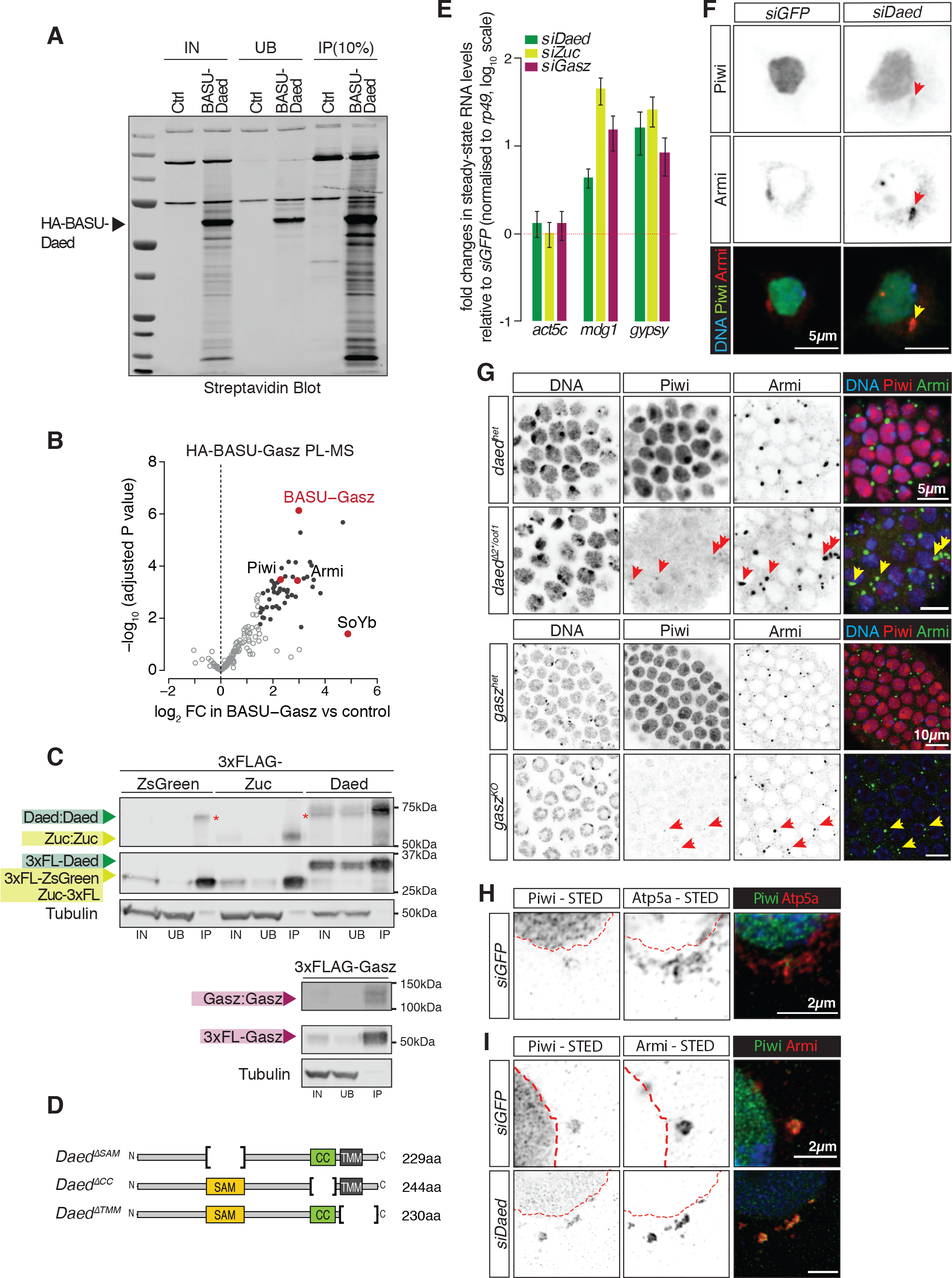
**(A)** Western blot showing pulldown of biotinylated proteins from OSCs expressing HA-BASU-Daed and control cells. Arrowhead indicates the size of the fusion protein. **(B)** Volcano plot showing enrichment and corresponding significance of biotinylated proteins identified via PL-MS from OSCs expressing BASU-Gasz against control (n=3). Black dots indicate proteins showing a log_2_FC > 1.5 and adjusted P value < 0.05. Highlighted in red are piRNA pathway factors. Full list of enriched proteins is provided in **Table S2**. **(C)** Western blot of FLAG-tag co-immunoprecipitates from lysates of chemically crosslinked S2 cells. Red asterisk indicates an unspecific band from anti-FLAG antibody. Arrowheads indicate the size of monomeric and dimeric proteins. **(D)** Schematic representation of Daed mutant constructs. **(E)** Fold changes in steady-state RNA levels of somatic transposons in OSC knockdown cells. Values are relative to *GFP* knockdown and normalised to *rp49*. Error bars indicate standard deviation (n=3). **(F-G)** Confocal images of Piwi and Armi in OSCs **(F)** and follicle cells of mutant flies **(G)**. Arrows point at sites of Piwi and Armi co-localization. **(H-I)** STED microscopy of Piwi and Atp5a **(H)** or Piwi and Armi **(I)** in OSCs. Scale bar, 2µm.

**Figure S5.**
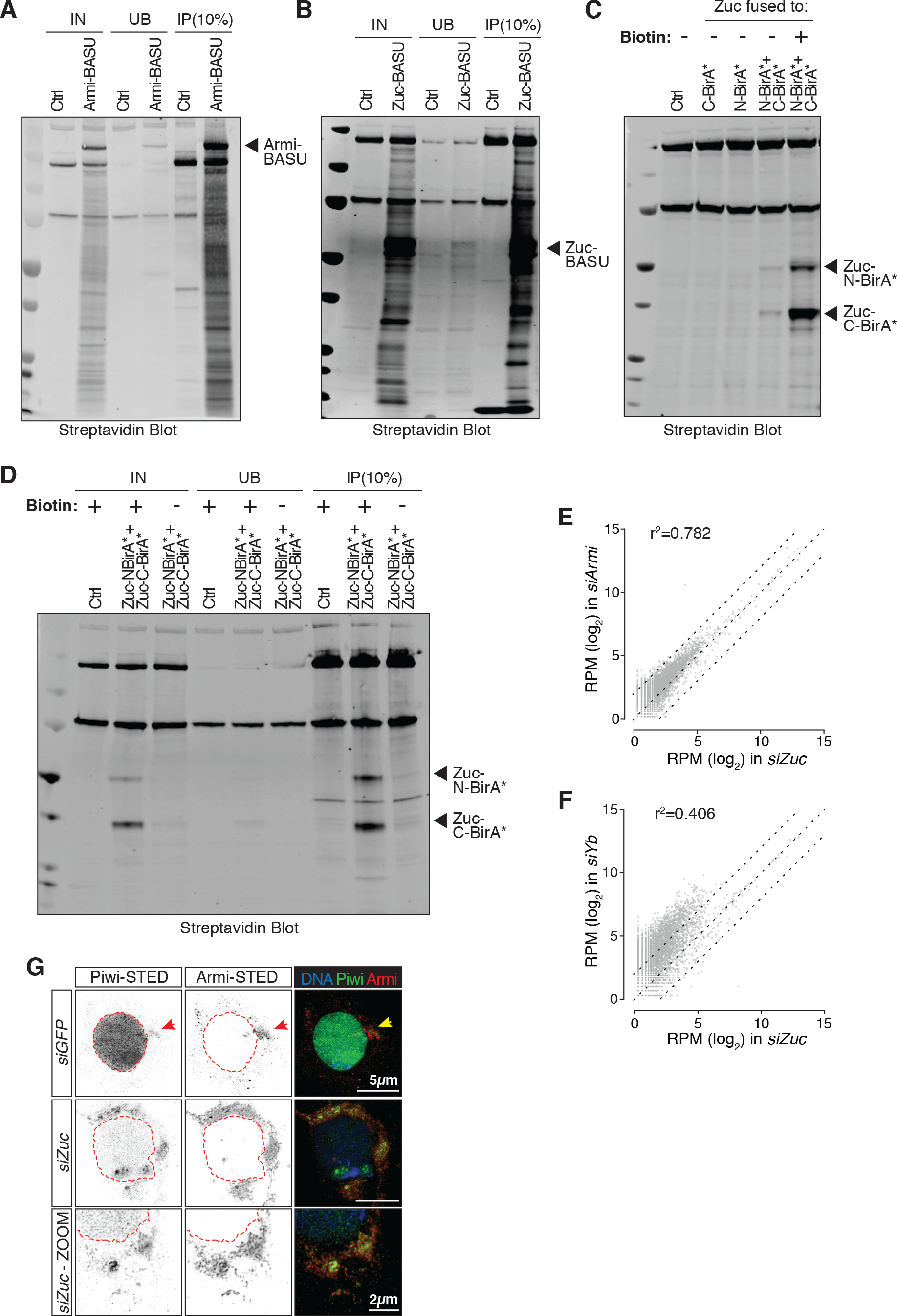
**(A,B,D)** Western blots showing pulldown of biotinylated proteins from OSCs. Arrowheads indicate the size of the fusion protein. **(C)** Western blot showing that expression of C-or N-BirA* fusions alone does not trigger protein biotinylation (lanes 2 and 3). Biotinylation can be readily detected only upon co-expression of both fusions and upon biotin supplementation (lane 5). **(E-F)** Scatter plots showing expression levels (normalised RPM) of genome-mapped small RNAs in OSCs upon indicated knockdown. **(G)** STED microscopy of Piwi and Armi in OSCs.

**Figure S6.**
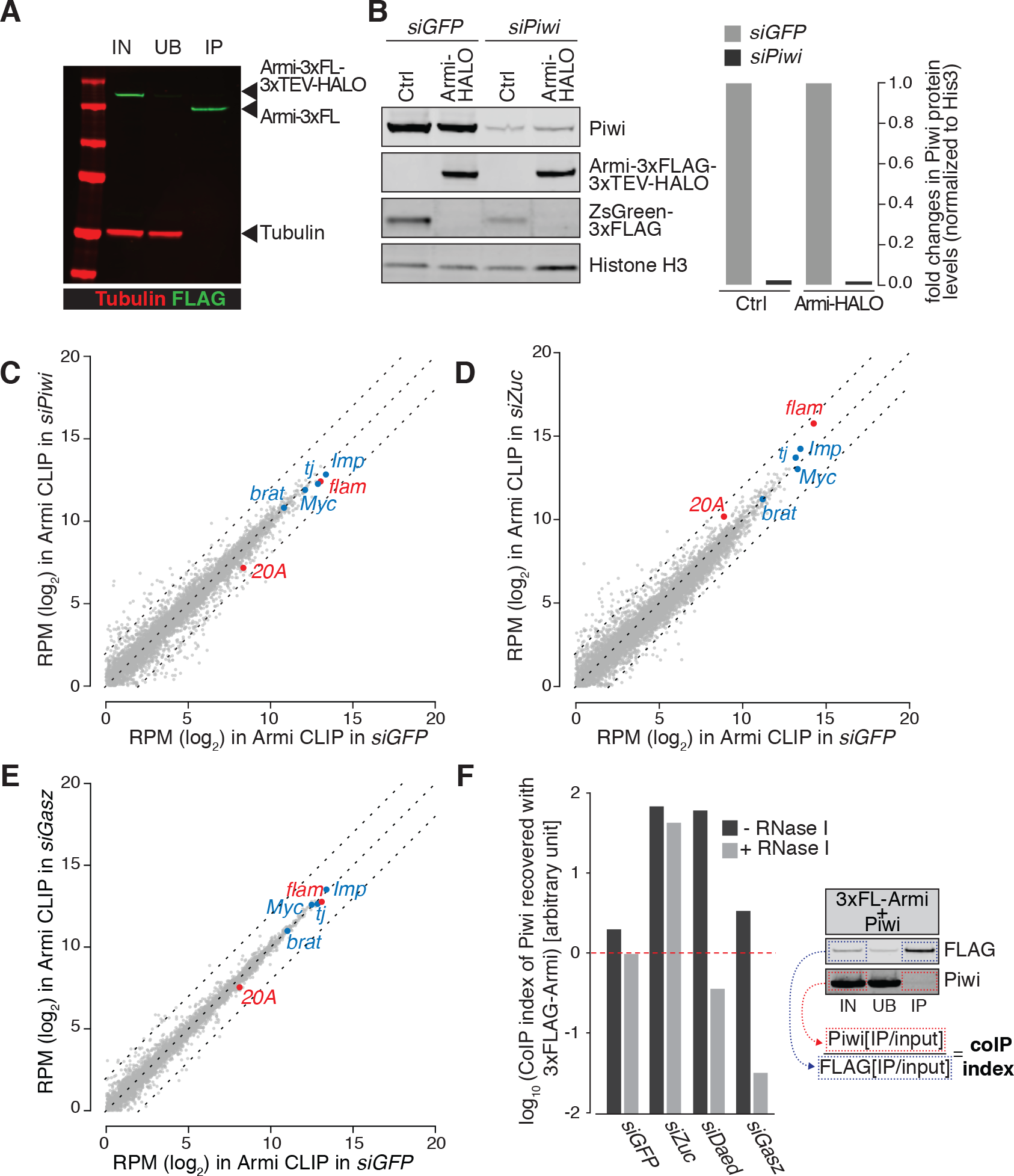
**(A)** Representative immunoprecipitation of the Armi-3×FLAG-3×TEV-HALO construct used for Armi CLIP-seq. **(B)** Western blot with relative quantification showing expression levels of Piwi in *siGFP* and *siPiwi* samples used for CLIP-seq in Fig.5A. **(C-E)** Scatter plots showing expression levels (RPM) of genes in Armi CLIP-seq upon indicated knockdowns. Somatic piRNA clusters expressed in OSCs are highlighted in red, selected protein coding genes producing piRNAs in blue. **(F)** CoIP index showing enrichment of Piwi in Armi-3×FLAG pulldown upon indicated knockdowns, relative to Figure 5E.

**Figure S7.**
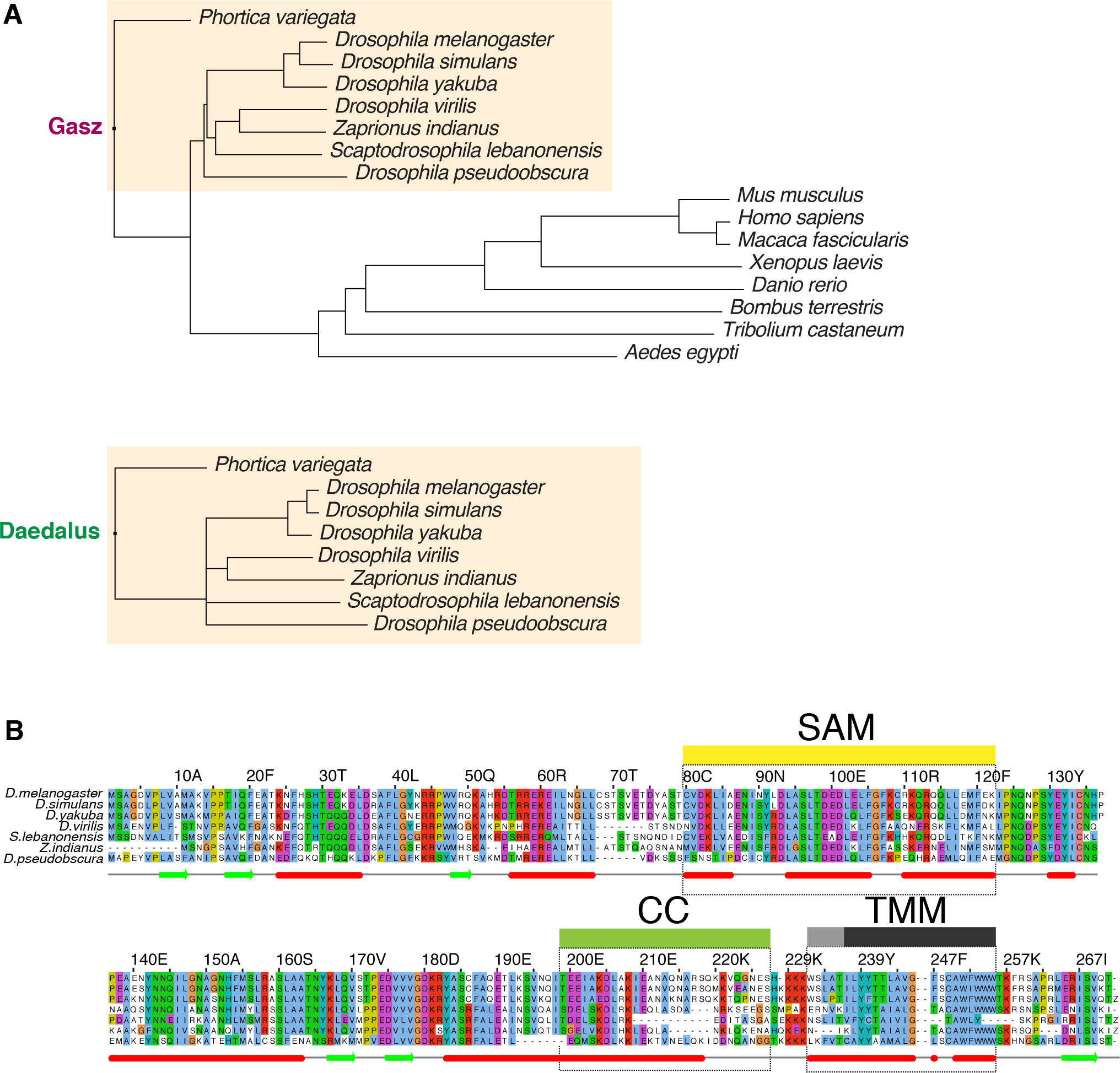
**(A)** Phylogenetic tree showing conservation of Gasz and Daed across species. Shaded in orange are *Drosophilidae* species. **(B)** Alignment of Daed amino acidic sequence from indicated species.

Table S1. Proteins detected in Streptavidin pulldowns from OSCs expressing BASU-Daed.

Table S2. Proteins detected in Streptavidin pulldowns from OSCs expressing BASU-Gasz.

Table S3. Proteins detected in Streptavidin pulldowns from OSCs expressing Armi-BASU.

Table S4. Proteins detected in Streptavidin pulldowns from OSCs expressing BASU-Zuc.

Table S5. Proteins detected in Streptavidin pulldowns from OSCs expressing Zuc split-BioID.

Table S6. Sequences of siRNAs and primers used in this study.

## AUTHOR CONTRIBUTIONS

M.M. performed all experiments with help from V.M., F.A.F. and B.C.. A.S. analysed the proteomics data, E.L.E. generated *daed* mutant flies and shRNA lines, E.K. contributed to the characterisation of their phenotype and J.W.E.S. generated *gasz* mutant flies. M.M., B.C. and G.J.H. designed the experiments, analysed and interpreted the data, and wrote the manuscript with input from the other authors.

## ACKNOWLEDGEMENTS

We thank Martin H. Fabry for help with computational analyses. We thank the CRUK Cambridge Institute Bioinformatics, Genomics, Microscopy, and Proteomics Core Facilities for support, in particular Kamal Kishore and Fadwa Joud. We thank the University of Cambridge Department of Genetics Fly Facility for microinjection services and fly stock generation. We thank the Vienna *Drosophila* Resource Center and the Bloomington Stock Center for fly stocks. We thank Mikiko Siomi for OSCs and anti-Armi antibody and Julius Brennecke for anti-Aub and anti-Ago3 antibodies. Research in the Hannon laboratory is supported by Cancer Research UK and by a Wellcome Trust Investigator award 110161/Z/15/Z. M.M. is supported by a Boehringer Ingelheim Fonds PhD fellowship.

## MATERIALS & METHODS

### Cell culture

Ovarian Somatic Cells (OSCs) were a gift from Mikiko Siomi and were cultured as described (Niki et al. 2006; Saito et al. 2009; Saito 2014). Knockdowns (all siRNA sequences are given in **Table S6**) and transfections in OSCs were carried out as previously described (Saito 2014). All constructs used in cells were expressed from the *Drosophila* act5c promoter. Full list of siRNAs is provided in **Table S5**. *Drosophila* Schneider 2 (S2) cells were purchased from Thermo Fisher Scientific and were grown at 26°C in Schneider media supplemented with 10% FBS. S2 cells were transfected using Effectene (Qiagen), according to manufacturer’s instructions.

### BASU Proximity Labelling and Mass Spectrometry

4×10^6^ OSCs were transfected with 20μg of plasmid expressing an HA-BASU fusion or HA-ZsGreen. After 48 hrs, the media was supplemented with 200μM Biotin for 1 hr. Cell pellets were lysed in 1.8 ml Lysis buffer (50mM Tris, pH 7.4, 500mM NaCl, 0.4% SDS, 1mM DTT, 2% Triton-100 with protease inhibitors) and sonicated using a Bioruptor Pico (Diagenode, 3× cycles 30 sec on / 30 sec off). Sonicated lysates were diluted 2× in 50mM Tris, pH 7.4 and cleared for 10 min at 16,500g. Following pre-clearing of the lysate with 100μl of Protein A/G Dynabeads (Thermo Fischer Scientific 10015D), biotinylated proteins were isolated by incubation with 200μl of Dynabeads (MyOne Streptavidin C1; Life Technologies) overnight at 4°C. The beads were washed 2× in 2% SDS, 2× in Wash Buffer 1 (0.1% deoxycholate, 1% Triton X-100, 500mM NaCl, 1mM EDTA, and 50mM 4-(2-hydroxyethyl)-1-piperazineethanesulfonic acid, pH 7.5), 2× with Wash Buffer 2 (250mM LiCl, 0.5% NP-40, 0.5% deoxycholate, 1mM EDTA, and 10mM Tris, pH 8), and 2× with 50mM Tris. Beads were rinsed twice with 100mM Ammonium Bicarbonate and submitted for Mass Spectrometry. HA-BASU-Daed pulldown was subjected to TMT-labelling followed by quantitative Mass Spectrometry on a nano-ESI Fusion Lumos mass spectrometer (Thermo Fisher Scientific). BASU-Gasz, Armi-BASU, Zuc-BASU and Zuc-SplitBioID pulldowns were analysed on a Q-Exactive HF mass spectrometer (Thermo Fisher Scientific). On bead Trypsin digestion and TMT chemical isobaric labelling were performed as described (Papachristou et al. 2018). Details on Mass Spectrometry analysis are available in Supplementary Materials and Methods.

### Split-BioID Proximity Labelling and Mass Spectrometry

4×10^6^ OSCs were transfected with 10μg of each plasmid expressing Zuc-CBirA*-6×His and Zuc-NBirA*-HA or 20µg of HA-ZsGreen. After 36 hrs, the growth media was supplemented overnight (~18 hrs) with 50mM Biotin. Harvesting and pulldown of biotinylated proteins were performed as stated above.

### Co-immunoprecipitation from cell lysates

S2 cells or OSCs were transfected with 3×FLAG- and HA-tagged constructs. After 48hrs, cells were lysed in 250μl of CoIP Lysis Buffer (Pierce) with Complete protease inhibitors (Roche). For cross-linking experiments, cell pellets were incubated with disuccinimidyl sulfoxide at 1mM final concentration (diluted in PBS) for 10’ at RT and 20’ at 4C followed by lysis in 50mM Tris, pH 7.4, 500mM NaCl, 0.4% SDS, 1mM dithiothreitol, 2% Triton-100 with protease inhibitors and sonication using a Bioruptor Pico (Diagenode, 3× cycles 30 sec on / 30 sec off). 200μg of proteins for each sample were diluted to 1 ml with CoIP Lysis Buffer and incubated with 30μl of anti-FLAG M2 Magnetic Beads (Sigma M8823) for 2 hrs at 4°C. The beads were washed 3×15 min in TBS with protease inhibitors, then resuspended in 2×NuPAGE LDS Sample Buffer (Thermo Fisher Scientific) and boiled for 3 min at 90°C to elute immunoprecipitated proteins.

### Western Blot

Images were acquired on an Odyssey CLx scanner (LiCor) using secondary antibodies (and/or Streptavidin; LiCor 925-32230) conjugated to infra-red dyes from LiCor. The following primary antibodies were used: anti-HA (ab9110), anti-FLAG (Sigma #F1804), anti-Piwi (Brennecke et al. 2007), anti-Atp5a (ab14748), anti-Tubulin (ab18251).

### OSCs immunostaining

Cells were plated one day in advance on Fibronectin-coated coverslips, fixed for 15 min in 4% PFA, permeabilized for 10 min in PBS, 0.2% Triton and blocked for 30 min in PBS, 0.1% Tween-20 (PBST) and 1% BSA. Primary antibodies were diluted 1:500 in PBST and 0.1% BSA and incubated overnight at 4°C. After 3×5 min washes in PBST, secondary antibodies were incubated for 1 hr at RT. After 3×5 min washes in PBST, DAPI was incubated for 10 min at RT and washed twice in PBST. Coverslips were mounted with ProLong Diamond Antifade Mountant (Thermo Fisher Scientific #P36961) and imaged on a Leica SP8 confocal microscope (100× Oil objective).

For STED, the same protocol was used with the following modifications: cells were plated on Fibronectin-coated 1.5H coverslips, blocking was for 1.5 hr in PBS, 0.1% Tween-20 (PBST) and 1% BSA. Primary and secondary antibodies were diluted 1:150 in PBST and 1% BSA. Coverslips were mounted using ProLong Glass Antifade Mountant (Thermo Fisher Scientific # P36982) and imaged on a Leica SP8 confocal microscope (100× Oil objective). The images were deconvoluted using Huygens Professional.

The following antibodies were used: anti-GFP (ab13970), anti-Atp5a (ab14748), anti-Piwi (Brennecke et al. 2007), anti-FLAG (Cell Signaling Technology 14793S), anti-HA tag (ab9111), anti-Armi (Saito et al. 2010).

### RNA isolation and qPCR analysis

Samples were lysed in 1ml Trizol and RNA was extracted according to manufacturer’s instruction. 1µg of total RNA was treated with DNAseI (Thermo Fisher Scientific), and reverse transcribed with the Superscript III First Strand Synthesis Kit (Thermo Fisher Scientific), using oligo(dT)_20_ primers. Real-time PCR (qPCR) experiments were performed with a QuantStudio Real-Time PCR Light Cycler (Thermo Fisher Scientific). Transposon levels were quantified using the ∆∆CT method (Livak and Schmittgen 2001), normalised to *rp49* and fold changes were calculated relative to the indicated controls. All oligonucleotide sequences are given in **Table S6**.

### Fly stocks and handling

All flies were kept at 25°C on standard cornmeal or propionic food. Flies carrying a BAC transgene expressing GFP-Gasz were generated by the Brennecke lab (VDRC # JB313277; (Handler et al. 2013). GFP-Zuc and GFP-CG10880 overexpression lines, shRNA-*daed*, *daed* mutant alleles (*CG10880^∆2*^* and *CG10880^oof1^*) and *gasz* mutant allele (*gasz^KO^)* were generated for this study (see below). Control *w^1118^* flies were a gift from the University of Cambridge Department of Genetics Fly Facility. For GLKD we used a stock containing a UAS::Dcr2 transgene and a nos::GAL4 driver (Czech et al. 2013) and shRNA lines from the Bloomington Drosophila Stock Center (BL35227) and Vienna Drosophila Resource Center (JB313133). Fertility of mutant females was scored by crossing ten freshly hatched females to five *w^1118^* males and counting the number of eggs laid in 12 hr periods and pupae that developed after 7 days.

### Generation of mutant and transgenic fly strains

Frameshift mutant alleles of *daed* were generated by injecting pCFD4 (Addgene plasmid # 49411; Port 2014) containing two gRNAs against CG10880 into embryos expressing vas-Cas9 (Bloomington stock 51323). *gasz^KO^* allele was generated by injecting a plasmid containing two gRNAs against gasz and a donor construct with 1kb homology arms flanking a 3×P3-RFP cassette into vas-Cas9 flies. shRNA against *daed* were cloned into pVALIUM20 (Ni et al. 2011), GFP-Daed and GFP-Zuc were cloned in an in-house generated transgenesis vector for phiC31-mediated integration and expressed under the *D.melanogaster* Ubiquitin promoter (pUBI). All plasmids were integrated into *attP40* sites on chromosome 2 (Stock 13-20). Microinjection and fly stock generation was carried out by the University of Cambridge Department of Genetics Fly Facility. Mutant flies were identified by genotyping PCRs and confirmed by Sanger sequencing.

### Ovary Immunostaining

Fly ovaries were dissected in ice-cold PBS, fixed for 15 min in 4% PFA at RT and permeabilized with 3×10min washes in PBS with 0.3% Triton (PBS-Tr). Samples were blocked in PBS-Tr with 1% BSA for 2 hrs at RT and incubated overnight at 4°C with primary antibodies in PBS-Tr and 1% BSA. After 3×10 min washes at RT in PBS-Tr, secondary antibodies were incubated overnight at 4°C in PBS-Tr and 1% BSA. After 4×10min washes in PBS-Tr at RT (DAPI was added during the third wash) and 2×5 min washes in PBS, samples were mounted with ProLong Diamond Antifade Mountant (Thermo Fisher Scientific #P36961) and imaged on a Leica SP8 confocal microscope. Images were deconvoluted using Huygens Professional. The following antibodies were used: anti-GFP (ab13970), anti-Atp5a (ab14748), anti-Piwi (Brennecke et al. 2007), anti-Aub (Senti et al. 2015), anti-Ago3 (Senti et al. 2015), anti-Armi (Saito et al. 2010).

### CLIP-seq

1×10^7^ OSCs were nucleofected first with 2µl of siRNA only and, 48 hrs later, with 2µl of siRNA and 5µg of the desired plasmid. 96hrs later, cells were crosslinked on ice with 150 mJ/cm^2^ at 254 nm. Cell pellets were lysed in 300µl of Lysis Buffer (50mM Tris-HCl pH 7.5, 150mM NaCl, 1% Triton^®^ X-100, 0.1% deoxycholate, Protease Inhibitor and RNasin Plus [1:500, Promega]),diluted to a final concentration of ~1µg/µl with 100mM Tris-HCl pH 7.5, 150mM NaCl and incubated with 200µl of Magne-HaloTag^®^ (Promega G7282) beads overnight at 4°C. Beads were washed 2× in Wash Buffer A (100mM Tris-HCl pH 7.5, 150mM NaCl, 0.05% IGEPAL^®^ CA-630), 3× in Wash Buffer B (PBS, 500mM NaCl, 0.1 % Triton X-100, RNasin Plus 1:2000), 3× in PBS, 0.1% Triton X-100, and rinsed in Wash Buffer A. Beads were resuspended in 100μl 1X ProTEV Buffer, 1mM DTT and RNasin Plus (1:50) and 25 units of ProTEV Plus Protease (Promega V6101) and incubated 2 hrs at 30°C. 15µL Proteinase K in 300µL PK/SDS buffer (100mM Tris, pH 7.5; 50mM NaCl; 1mM EDTA; 0.2% SDS) were added to the eluate and incubated 1hr at 50°C. RNA was isolated with Phenol-Chlorophorm and library preparation was carried out with the SMARTer^®^Stranded RNAseq kit (Takara Bio 634839), according to manufacturer’s instructions. CLIP-seq libraries were sequenced on an Illumina HiSeq 4000 (Illumina).

### Small RNA-seq library preparation

Small RNA libraries were generated as described previously with slight modifications (McGinn and Czech 2014). Briefly, 18-to 29-nt-long small RNAs were purified by PAGE from 15µg of total RNA from ovaries or OSCs. Next, the 3’ adapter (containing 4 random nucleotides at the 5’ end (Jayaprakash et al. 2011) was ligated using T4 RNA ligase 2, truncated KQ (NEB). Following recovery of the products by PAGE purification, the 5’ adapter (containing 4 random nucleotides at the 3’ end) was ligated to the small RNAs using T4 RNA ligase (Ambion). Small RNAs containing both adapters were recovered by PAGE purification, reverse transcribed and PCR amplified. Libraries were sequenced on an Illumina HiSeq 4000. All adapter sequences are given in **Table S6**.

### CLIP-seq and small RNA-seq analysis

Details on sequencing analysis are available in Supplementary Materials and Methods.

### Data availability

Raw data from proteomics and high-throughput sequencing experiments are available on PRIDE (<u>PXD013417</u>, <u>PXD013405</u>, <u>PXD013404</u>, <u>PXD013403</u>) and GEO (<u>GSE129321</u>).

