## Supplementary Materials and Methods for "Daedalus and Gasz recruit Armitage to mitochondria, bringing piRNA precursors to the biogenesis machinery"

### Mass Spectrometry analysis

Spectral .raw files from PL-MS of BASU-Daed were processed with the SequestHT search engine on Thermo Scientific™ Proteome Discoverer™ 2.1. Data was searched against a custom FlyBase database (*"dmel-all-translation-r6.24"*) at a 1% spectrum level FDR criteria using Percolator (University of Washington). MS1 mass tolerance was constrained to 20 ppm and the fragment ion mass tolerance was set to 0.5 Da. TMT tags on lysine residues and peptide N termini (+229.163 Da) were set as static modifications. Oxidation of methionine residues (+15.995 Da), deamidation (+0.984) of asparagine and glutamine residues, and biotinylation of lysines and protein N-terminus (+226.078) were included as dynamic modifications. For TMT-based reporter ion quantitation, we extracted the signal-to-noise ratio for each TMT channel. Parsimony principle was applied for protein grouping and the level of confidence for peptide identifications was estimated using the Percolator node with decoy database search. Strict FDR was set at q-value < 0.01. Downstream data analysis was performed on R using the qPLEXanalyzer package (https://doi.org/10.5281/zenodo.1237825) as described (Papachristou et al. 2018). Only proteins with more than one unique peptide were plotted.

Spectral .raw files from PL-MS of BASU-Gasz, Armi-BASU, Zuc-BASU, Zuc Split-BioID were processed with the SequestHT search engine on Thermo ScientificTM Proteome Discoverer™ 2.2. The node for SequestHT included the same parameters as above (except Fragment Mass Tolerance set to 0.02 Da) and modifications. The Precursor Ion Quantifier node (Minora Feature Detector) included a Minimum Trace Length of 5, Max. ΔRT of Isotope Pattern 0.2 minutes. For calculation of Precursor ion intensities, Feature mapper was set True for RT alignment (mass tolerance of 10ppm). Precursor abundance was quantified based on intensity and the level of confidence for peptide identifications was estimated using the Percolator node with a Strict FDR at q-value < 0.01. Analysis of label-free quantification protein intensity data was carried out in R (v 3.5.1) using the *qPLEXanalyzer* package (v 1.0.3) (Papachristou et al. 2018). Peptides for which all control (ZsGreen) samples lacked measurements or for which more than 1 target protein sample lacked measurements were discarded. Remaining missing values were then imputed using the nearest neighbour averaging (knn) imputation method provided in the R package *MSnbase* (v 2.8.3) (Gatto and

Lilley 2012). Differential analysis was carried out by linear modelling using *limma* based methods provided by the *qPLEXanalyzer* package. Multiple testing correction of *p-values* was applied using the Benjamini & Yekutieli method to control FDR (Benjamini et al. 2001). Only proteins with more than one unique peptide were plotted.

**CLIP-seq and small RNA-seq analysis**

Raw fastq files generated by Illumina sequencing were analysed by a pipeline developed in-house. In short, for CLIP-seq the first 5 bases of each 50 bp read were removed using fastx trimmer (http://hannonlab.cshl.edu/fastx_toolkit/). After removal of rRNA mapped reads, high-quality reads were aligned to the *Drosophila melanogaster* genome release 6 (dm6; downloaded from Flybase) using STAR (Dobin et al. 2013). For transposon-wide analysis, genome multi-mapping reads were randomly assigned to one location using option '--outFilterMultimapNmax 1000 --outMultimapperOrder Random' and non-mapping reads were removed. Alignment files were then converted back to fastq format with samtools (Li et al. 2009) and re-aligned to the transposon consensus sequences allowing multi-mappers that were assigned to a random position. Generated bam alignment files were indexed using samtools index. For genome-wide analyses, multi-mapping reads were removed to ensure unique locations of reads. Normalization was achieved by calculating rpm (reads per million) using the deepTools2 bamCoverage function with 10 bp bin sizes (Ramirez et al. 2016). Reads mapping to genes were counted with htseq (Anders et al. 2015) and transposon derived reads were calculated using a custom script. Differential expression analysis was performed using custom built R scripts. Coordinates used for the piRNA clusters are chrX:21631891-22282863 (*flam*) and chrX:21520428-21556793 (*20A*).

For small RNA-seq, adapters were clipped from raw fastq files with fastx_clipper (adapter sequence AGATCGGAAGAGCACACGTCTGAACTCCAGTCA) keeping only reads with at least 23 bp length. Then the first and last 4 bases were trimmed using seqtk (https://github.com/lh3/seqtk). After removal of cloning markers and 2S rRNA mapped reads, alignment was performed as described above and normalised to miRNA reads in the control library (set to rpm). Only high-quality small RNA reads with a length between 23 and 29 bp were used for further analysis of small RNA profiles. piRNA distribution was calculated and plotted in R. The ping-pong signature was calculated using piPipes (Han et al. 2015).
